# PIF4 promotes water use efficiency during fluctuating light and drought resistance in rice

**DOI:** 10.1101/2023.03.02.530909

**Authors:** Sushuang Liu, Jemaa Essemine, Yanmin Liu, Chundong Liu, Feixue Zhang, Zhan Xu, Mingnan Qu

## Abstract

Molecular mechanism of intrinsic water use efficiency (iWUE) during fluctuating light (FL) was rarely understood. In this study, we investigated five parameters of iWUE under FL in 200 Minicore rice accessions. Among them, a novel trait, WUE_FL_ (averaged iWUE during FL) has highest SNP heritability in these parameters. GWAS identifies six candidate genes, and *PIF4* is highly expressed in high iWUE_FL_ rice subgroup. Nine SNPs were significantly associated with iWUE_FL_, and v3 SNP located at -1,075 bp of *PIF4* promoter shows highest sensitives to light. Deletion of v3 in a rice cultivar, WYG7 (PIF4^v3m^) leads to ∼20% reduction in iWUE_FL_, and overexpressing PIF4 causes 25% increase in iWUE_FL_ under DS. There are 85% reduction in adenosine 3’,5’-diphosphate (PAP) amounts together with 73% increase in *SAL1* gene abundance in PIF4^v3m^ than WYG7. PIF4 transcriptionally repress and activate *SAL1* and *NHX1*, respectively, through binding to G-box motifs of the two genes, which leads to 16% reduction and 5% increase in iWUE_FL_ in co-overexpression rice lines of *PIF4*-*SAL1* and *PIF4*-*NHX1*, respectively, relative to *PIF4*-OE under DS. We proposed that PIF4 promotes iWUE_FL_ and stomatal adjustment via targeting the G-box motif of *SAL1* and *NHX1* genes during FL, eventually facilitating to drought resistance.

## Introduction

Water for agricultural use becomes increasingly scarce due to climate change and rapid industrialization and urbanization (Yan *et al*., 2015). By the year 2025, irrigated rice of 15–20 million hectares in Asia will suffer water scarcity (Tuong & Bouman, 2003). Farmers are facing a challenge to produce more rice per unit land with limited water in order to meet the food demand of the growing population. Therefore, it is urgent to breed new rice cultivars with greater yield potential with consideration of improved water use efficiency (WUE) (Horie *et al*., 2004).

There are several parameters that used to be indicators of WUE across leaf to whole-plant levels, include δ^13^C, thermal imaging and whole-plant WUE. δ^13^C is proved to be negatively related to the ratio of *A* to stomatal conductance (*g*_s_) through *p*_i_/*p*_a_, the ratio of intercellular and ambient CO_2_ partial pressures (Farquhar & Richards, 1984). Thermal imaging has been extensively used to monitor stomatal closure and WUE (McAusland *et al*., 2013; Vialet-Chabrand & Lawson, 2019). Whole-plant WUE can be expressed as the ratio of biomass or grain production to the amount of water consumed (Medrano *et al*., 2015). These parameters were proved to be as important indicators used in drought targeted breeding.

At leaf levels, there are two types of WUE, i.e., intrinsic and instantaneous water use efficiency, calculated by the ratio of *A* to *g*_s_, and the ratio of *A* to transpiration (T), respectively. The two leaf level WUE parameters were extensively used in various studies (Gornall & Guy, 2007; Broeckx *et al*., 2014; Medrano *et al*., 2015). This is thanks of the application of portable equipment for facilitating the simultaneous measurement of *A*, *g*_s_ and T. However, the use of these WUE parameters remains debatable, because it was reported to show poor relationship with WUE parameters at whole-plant level (Medrano *et al*., 2015). This is because light in daily is always changed, therefore changing light regime should be considered simultaneously.

Fluctuating light (FL) is an importantly environmental factor in iWUE through stomatal regulation (McAusland *et al*., 2016). There is large nature variation in stomatal kinetics and hence iWUE reported in different species (Lawson *et al*., 2012). Even within one species, our previous studies suggest that large genetic variation was also observed (Qu *et al*., 2016). Many regulators mediating stomatal kinetics during FL were reported, such as *PATROL1* (Hashimoto-Sugimoto *et al*., 2013), *SLAC1* (Yamori *et al*., 2020), *NHX1* (Qu *et al*., 2020). Genome-wide association study (GWAS) emerges as an important tool in gene mining, but very less studies were reported about gene regulation of iWUE under FL using GWAS (Pignon *et al*., 2021).

In this study, we investigated the natural variation of iWUE dynamics during FL treatment in Minicore rice population consisting of 200 rice accessions (Agrama *et al*., 2009). It is proved as an ideal germplasm since it has a suitable population size and sufficient genetic diversity for gene mining on photosynthetic and nitrogen use related traits (Qu *et al*., 2020; Liu *et al*., 2021). Based on genome-wide association study, overlapped lead SNPs among different iWUE related traits were identified, and the function of one promising candidate gene, PIF4 was validated by genetic, molecular biological, and biochemical techniques. This study provides a deep insight into the genetic mechanism of PIF4 regulating iWUE dynamic through coordinating with SAL1 and NHX1 during FL in rice population, which helps to guide the molecular selection breeding towards improved grain production and water use.

## Materials and Methods

### Plant growth conditions and experimental set-up

In total, Minicore rice population consists of 200 accessions derived from 97 countries were used in this study. The population contains 108 accessions from *indica* (IND, AUS), 73 accesions from *japonica* (TEJ, TRJ and ARO) and 19 accessions with admix type. The population contains six subpopulations *indica* (35.4%), *aus* (18.7%), *tropical japonica* (18.2%), *temperate japonica* (15.2%), *aromatic* (3.0%) and their admixtures (9.6%) (Agrama *et al*., 2009; Li *et al*., 2010). The population were grown in Nov 2020 in paddy field in Hainan (110.0375E, 18.5060N), China. To decrease the boundary effect, 49 plants were planted (7×7) for each rice accession, with 15 cm between plants within each row and 20 cm between rows. The rice fields were managed according to standard local agronomic practice with the following fertilizer application guideline: 48 kg N ha^-1^, 120 kg P_2_O_5_ ha^-1^, and 100 kg K_2_O ha^-1^ as the basal fertilizer, and additional 86 kg N ha^-1^ at the tillering stage and 28 kg N ha^-1^ at the booting stage. For moderate drought stress (DS) treatments, plants were exposed to limited irrigation from 30 to 50 days after germination (DAG). The soil humidity was maintained ranging of 25∼40% with monitored by a soil moisture meter (Hansatech Instruments Ltd, UK).

### Generation of transgenic plants and near isogenic lines

To study the function of *PIF4* (*LOC_Os03g56950*) based on genome-wide association study, a *PIF4* mutant (*PIF4^v3m^*) was generated with four nucleotide knockout including a SNP using CRISPR/CAS9 by the Biogle Company (Hangzhou, China). The sgRNA was inserted into the BGK032-DSG vector containing Cas9, which was introduced into an *Agrobacteruim tumefaciens* strain EHA105 and transformed into the wild-type Wuyungeng 7 (WT, WYG7). The homozygous lines were sequenced based on the gene specific primers listed in Table S1.

To construct the overexpression vector for *PIF4, SAL1* (inositol polyphosphate 1-phosphatase, LOC_Os07g37220) and *NHX1* (vacuolar sodium/proton antiporter, LOC_Os07g47100), its cDNA (primer sequences are listed in Table S1) was amplified from WYG7 and restriction sites (*Bam*HI and *Sac*I) were added and transferred into pCAMBIA 1301 plasmid backbone. This backbone includes a GFP tag (Youbio, China, VT1842) and hygromycin B phosphotransferase (*HPT*) gene driven by 35S promoter (cauliflower mosaic virus). Eventually, the synthesized 35S:*SAL1*-GFP and 35S:*NHX1*-GFP constructs were transformed into rice following the earlier reported protocol (Shan *et al*., 2014) to generate different over-expression lines (OE). Co-overexpression lines of two genes (*PIF4*-*SAL1* and *PIF4*-*NHX1*) were obtained by crossing the OE lines of *PIF4* with other OE lines of *SAL1* and *NHX1*, respectively.

The T_1_ generation of different OE lines were planted in a paddy field in Hainan (E110°02′, N18°48′), China, in May 2021. The primers used to detect the positive lines were also listed in Table S1. Three homozygous OE lines with highest expression of *PIF4* gene from the T_3_ generation were selected for the drought and FL treatment experiments. Primers used for detecting the *HPT* expression level are also listed in Table S1.

### Kinetics of iWUE measurements during fluctuating light

In this study, plants at ∼50 DAG exposed to 20 d DS were first incubated in a 12h with 30°C/25°C and 60∼70% for air temperature and humidity, respectively. To ensure stomatal adaption, a 2 h high light treatment (1500 μmol m^-2^s^-1^ PPFD) was applied before iWUE kinetics measurements (Qu *et al*., 2016). Four portable photosynthesis systems LICOR 6800 (Li-COR, Inc.) were used to measure kinetics of gas exchange parameters driven by a FL treatment from 8:30 am to 16:30 o’clock. The FL treatment (40 min in total) was started by 10 min high light (HL), followed by 25 min low light (LL) and restored 5 min as mentioned above. The FL treatment was controlled in leaf cuvette of LICOR 6800. These gas exchange parameters include net photosynthetic rates (*A*), stomatal conductance (*g*_s_), intrinsic water use efficiency (iWUE) under HL (1500 μmol m^-2^s^-1^ PPFD). Fully-expanded leaves were chosen with 400 μmol mol^-1^ CO_2_ concentration and 500 μmol mol s^-1^ flow rates in leaf cuvette. Four biological replicates were conducted for each rice accession.

### The calculation of iWUE parameters

There are five time-points during light switch according to dynamics of *A*, *g*_s_ and iWUE as observed previously (Qu *et al*., 2016). First, iWUE sharply decreased when light switched from high to low light from time-point *i* to *j*. Second, iWUE increases linearly due to *g*_s_ deep decrease from time-point *j* to *k*. Third, iWUE starts slightly decrease due to *g*_s_ recovery driven by its plasticity from time-point *k* to *p*. Last, light was restored and iWUE was rapidly increased from time-point *p* to *f*. According to different phases, five iWUE related parameters were calculations as following equations:

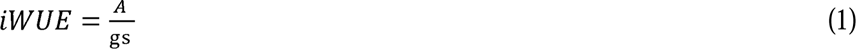

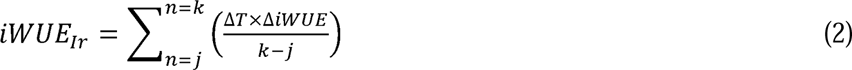

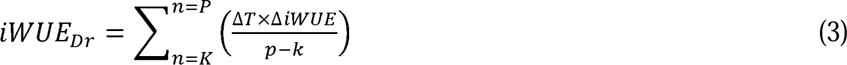

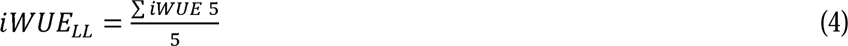

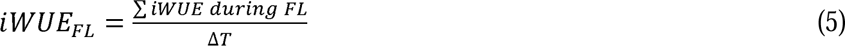

where Δ*T*: time interval from each point during FL; iWUE_Ir_: slope of iWUE during linear increase phase of FL (from *j* to *k*); iWUE_Dr_: slope of iWUE when third stage of dropping light (from *k* to *p*); iWUE_LL_: steady-state iWUE of five data points at end of low light phase when iWUE appears to be plateau; iWUE_FL_: totally averaged iWUE during FL. For stomatal closure speed data analysis, we fitted the stomatal dynamics data with a first order exponential decay curve using MATLAB software, version R2010a (Mathworks Inc., Natick, MA, USA) to estimate τ_cl_ as reported in (Vico *et al*., 2011).

### Fluctuating grown light treatment

To mimic the effects of FL on iWUE and drought resistance, 10 rice accessions from Minicore population consisting of 5 accessions with extremely high iWUE_FL_ and 5 accessions with extremely low iWUE_FL_ were grown in the phytotron. Plants were grown in the pot (12L volume) with four plants sown in one pot. The air temperature, humidity and photoperiod in phytotron maintain at 30°C/25°C, 60∼70% and 12h/12h, respectively. To ensure consistency of nutrition supply, a commercial peat soil (Pindstrup Substrate no. 4, Pindstrup Horticulture Ltd, Shanghai, China) was used. Plants were treated with limited irrigation as mentioned above in Minicore population. In addtion, plants were simultaneously exposed to a FL treatment (40 min in total) as mentioned above. The FL treatment was applied by a light density controller (FL053C, Zhongkang Omics Corp. Ltd. China). The light intensity was estimated at 30 cm distance from LED light source via light meter (Quantitherm light meter, Hansatech, UK). The FL treatment was stopped at 50 DAG until kinetics of iWUE measurements started. Thermal imaging of plants was used to estimate the changes of leaf temperature due to transpiration with a thermal imager equipped with FLIR E40bx IR camera (Teledyne FLIR LLC, UK).

### δ13C determination and stomatal aperture determinations

An elemental analyzer-isotope ratio mass spectrometer (EA-IRMS, EA model of Flash2000 and IRMS model of MAT253) (Thermo Fisher) was used to determine δ^13^C. The measurement of plant δ^13^C value was based on the PDB (Pee Dee Belemnite) as previously documented (Farquhar & Richards, 1984). For stomatal aperture determinations, top fully-expanded leaves of each plant were selected to cut and stored in formalin-acetic-alcohol (FAA) fixative solution for future use. The detailed process of stomatal observation was followed as previously documented (Chen *et al*., 2020). TM-1000 desktop scanning electron microscope was used to observe stomatal characteristics on adaxial leaf surface.

### Genome-wide association study

A genome wide association study (GWAS) was performed on the genotype dataset generated from low coverage genome sequencing published (Wang *et al*., 2016), using a mixed linear model implemented in GEMMA (version 0.98.5) (Zhou & Stephens, 2012). A relatedness matrix and the first four principal components from the PCA were used for controlling population structure. To control for inflation in the genome-wide association tests, we obtained the genome-wide significance level by performing permutations: the phenotypes were permuted 200 times. Both Manhattan plot and QQ plot for GWAS were generated using the R package qqman (Turner, 2014). The Locus Zoom plot was generated by R script from Github (https://github.com/statgen/ locuszoom-standalone).

### Haplotype analysis

A linkage disequilibrium (LD) analysis was also conducted with the Haploview software (version 4.2) to investigate the relatedness degree of the candidate genes to the lead SNP. The LD blocks were generated when the upper 95% confidence bounds of *r^2^* value exceeds 0.98 and the lower bounds exceeds 0.70 (Gabriel *et al*., 2002). The genes identified in the LD block were selected as potential candidate genes that might control iWUE related parameters.

### Estimation of SNP heritability

The SNP heritability (*h*^2^_SNP_) represents the proportion of phenotypic variance explained by SNPs (Speed *et al*., 2017). We estimated the *h*^2^_SNP_ of the iWUE related parameters using the GCTA software (version 1.11.2 beta) based on the Restricted Maximum Likelihood method. A Log Likelihood Test (LRT) was used to calculate the *P*-value of SNP heritability according to the GCTA software manual.

### GO and KEGG analysis

For Gene Ontologies (GO) annotation on the candidate genes corresponded to the overlapped SNPs among different iWUE parameters, we used an in-house Perl script UniProtKB GOA file (ftp.ebi.ac.uk/pub/databases/GO/goa). KOBAS (KEGG Orthology Based Annotation System, v2.0) was applied to identify reprogrammed biochemical pathways of each pathway as previously documented (Xie *et al*., 2011).

### RNA extraction and qRT-PCR analysis

To further identify the candidate genes responsible for both iWUE and iWUE_FL_, we performed a qRT-PCR analysis in different rice accessions. Top fully-expanded leaves from each plant exposed to DS coincident with FL were sampled for qRT-PCR analysis. Total mRNA was extracted from the leaves in WYG7 exposed to FL via TRIzol Reagent (Invitrogen) Genomic DNA was removed with DNase I (Takara). cDNA was then reverse transcribed via SuperScript VILO cDNA Synthesis Kit (Invitrogen Life Technologies). The qRT-PCR analysis was performed using SYBR Green PCR Master Mix (Applied Biosystems, USA, 4309155) via a Real-Time PCR System (ABI StepOnePlus, Applied Biosystems, USA). Primers for qRT-PCR were designed using Primer Prime Plus 5 Software Version 3.0 (Applied Biosystems, USA). For internal reference, we used *Actin1* gene (LOC_Os03g50885). Relative expression of gene against *Actin1* was calculated as: 2^−ΔΔCT^ (ΔCT = CT, gene of interest^−CT^), as described earlier (Livak & Schmittgen, 2001). The ratio of (dark / HL) for the expression levels of each gene was used to indicate light response. Three biological replicates were used. The primers used for determining iWUE and iWUE_FL_ responsive gene expression levels are listed in Table S1.

### Determination of PIF4 subcellular localization

We used transient transformation in tobacco leaves to study the PIF4 subcellular localization. A GFP fusion vector *p35S::PIF4-GFP* was constructed using pCAMBIA1300 backbone. A known nuclear localized bZIP transcription factor EmBP1 was used as positive control as reported previously (Perveen *et al*., 2020). The construct was transformed into tobacco leaves through agro-infiltration method (INZÉ *et al*., 2012). Briefly, a GFP-protein fusion construct was transfected into the *Agrobacterium tumefaciens* strain C58C1 (WeiDi Biotech, China, AC1110). The constructs were then transiently expressed in leaf epidermal cells of a 5-week-old tobacco (*Nicotiana benthamiana*) by *Agrobacterium*-mediated leaf infiltration(Sparkes *et al*., 2006). Green fluorescence of the GFP fusion protein was detected using a confocal laser scanning microscope (Zeiss LSM 700, Germany) with a Fluar 10X/0.50M27 objective lens and an SP640 filter. GFP and chlorophyll autofluorescence (Chl) were excited at 488 nm and signals were collected at 660–736 nm (Chl) and 495–515 nm (GFP).

### Luciferase activity assay in rice protoplasts

For analysis of the *PIF4* promoter in response to FL, approximately 2.0-kb DNA fragments upstream of *PIF4* coding region were amplified from WYG7 and inserted into pGreenII 0800-LUC vector to generate proPIF4-L:LUC and proPIF4-H:LUC, respectively. Six variants were mutated on the basis of proPIF4-L:LUC using Q5 site-directed mutagenesis kit (NEB, E0552S) according to the manufacturer’s protocol. The primers used for PCR amplification and mutation are listed in Table S1. One-month-old rice seedlings grown in high light regime (∼1500 μmol m^-2^s^-1^ PPFD). Before protoplast extraction, plants were first exposed to dark for 24 h, and leaf samples were collected for protoplasts. All the vectors were transformed into protoplasts, and each transformation product was divided into two portions, exposing dark and high light for 1h to generate dark and high-light conditions, respectively. Afterwards, all the products were incubated in W5 solution for 4–6 h at 28 °C. Activities of firefly luciferase (LUC) and *Renilla* luciferase (REN) were examined using a Dual-Luciferase Reporter Assay System kit (Promega, E1960). LUC/REN was calculated as the relative activity and the ratio of dark / HL was used to indicate light response. For each vector, three replicates were used to evaluate the light response.

### mRNA extraction and library preparation

Total RNA was extracted using TRIzol reagent according to the manufacturer’s instructions (Invitrogen, Carlsbad, CA). RNA degradation and contamination was monitored on 1% agarose gels, and purity was checked using the Nano-Photometer spectrophotometer (IMPLEN, CA, USA). RNA integrity was assessed using the RNA Nano 6000 Assay Kit of the Agilent Bioanalyzer 2100 system (Agilent Technologies, CA, USA). A total amount of 1.5 μg RNA per sample was used as input material for the RNA sample preparations. Sequencing libraries were generated using NEB Next Ultra RNA Library Prep Kit for Illumina (NEB, USA) as reported previously (Jiang *et al*., 2020). After cluster generation, the library preparations were sequenced on an Illumina Hiseq 4000 platform and 150 bp paired-end reads were generated.

### Read mapping and differentially expressed analysis

The quality of RNA-seq data from WYG7 and PIF4^v3m^ was assessed by FastQC software. After generating the genome index, the clean RNA-seq reads were aligned by STAR (Dobin *et al*., 2013) with ‘—quantMode GeneCounts’ option to count number of reads per gene. Quantification of genes and isoforms was performed using cufflinks version 2.2.1. To identify differentially expressed genes (DEGs) between WYG7 and PIF4^v3m^ under fluctuating light coincident with moderate drought stress, a fragment per kilobase of transcript per million mapped reads (FPKM) method was applied to calculate the transcript abundance. Notably, DEGs were determined by the R package ‘DESeq2’ (Love *et al*., 2014) with the reads counts reported by STAR (Dobin *et al*., 2013). Only genes with the adjusted *P*-value <0.05 were considered as DEGs. To reduce transcription noise, each isoform/gene was included for analysis only if its FPKM values was > 0.01, a value was chosen based on gene coverage saturation analysis as described earlier (Zhou *et al*., 2022) .

### Electrophoresis mobility shift assay (EMSA) experiments

Full-lengths of PIF4 were amplified and cloned into the pET51b vector as documented previously (Perveen *et al*., 2020), while GFP protein was used as negative control. Strep–PIF4 recombined proteins were expressed in the Escherichia coli BL21 (DE3) strain and then purified using amylose resin (NEB, E8021S). DNA probes of NHX1 and SAL1 were synthesized and labeled using the Biotin 3′ End DNA Labeling Kit (Thermo Fisher Scientific, 89818). The EMSA was performed using the LightShift Chemiluminescent EMSA kit (Thermo Fisher Scientific, 20148). The probe sequences are shown in Table S1.

### LC-MS/MS metabolism analysis and metabolites identification

A non-targeted metabolic profiling in the leaves of both WYG7 and PIF4^v3m^ exposed to DS was prepared based on the LC-MS/MS (Q Exactive, Thermo Scientific) procedure. Shortly, ∼2.5 mg leaf samples collected from WYG7 and PIF4^v3m^ were sampled in 2 ml Eppendorf tube containing pre-cooled metal beads, then immediately stored in liquid nitrogen. The samples were pretreated as previously reported (Li *et al*., 2020). The metabolomic analysis was performed using metabolon software (Durham, NC, USA). Regarding the metabolic compounds identification of each sample, the mass spectra with the entries of the mass spectra libraries NIST02 and the Golm metabolome database were considered (http://csbdb.mpimp-golm.mpg.de/csbdb/gmd/gmd.html).

### PAP content determinations

Total adenosines were extracted with 0.1 M HCl, derivatization with chloroacetaldehyde and quantified fluorometrically after HPLC fractionation as previously described (Bürstenbinder *et al*., 2007). 3′-phosphoadenosine 5′-phosphate (PAP) quantification was performed by integrating the HPLC peak area and converting these to pmol units using standard curves of 1, 5 and 10 pmol standard.

### Western blots analysis

To compare the protein expression levels of PIF4 and SAL1 in overexpression lines of two genes, western blot experiments were performed. Briefly, about 5 µg total protein of crude extracts from leaves, and protein concentration was determined using BCA protein concentration assay kit (Beyotime, P0010). The extracts were then loaded and separated on a 12% SDS–PAGE gel, then the gel was stained with Coomassie brilliant blue (CBB), or transferred to a nitrocellulose membrane for western blotting analysis according to the methods as described (He & Mi, 2016) and signals were detected using a Pierce ECL Plus Kit (Thermo Scientific, USA), and visualized with a luminescent image analyzer (Tanon-5200, Tanon). The antibody of PIF4 (cat#AS163955) and SAL1 (cat#AS07256) against rabbit were purchased from Agrisera (USA) with dilution 1:10,000.

### Yeast-one hybrid assays

To detect whether PIF4 binds to the G-box CACGTG-motif present in the *NHX1* and *SAL1* promoters in yeast, Yeast one-hybrid (Y1H) assays were performed according to the manufacturer’s instruction (Clontech, CA, USA). The coding region of PIF4 was amplified and cloned into the yeast GAL4 activation domain (GAL4 AD) of pGADT7-Rec2 (Clontech, CA, USA) prey vector. The promoters (∼1.5k-bp upstream of ATG) of *SAL1* and *NHX1*, harboring G-box CACGTG-motifs, were amplified and cloned into pAbAi (Clontech, CA, USA) to generate the bait vectors (Pro-*SAL1*-AbAi and Pro-*NHX1*-AbAi). In addition, the two motifs were both mutated by PCR and also ligated into pAbAi. The prey vectors were co-transformed with bait vector into Y1H Gold yeast strain. The concentration of positively transformed yeast cells that harbor different combinations of the bait and prey vectors were adjusted to OD600 ≈ 1.0 and serially diluted 1/10, 1/100, 1/1000 in sterile ddH_2_O. The serial dilution of transformed yeast cells was cultured on SD/−Leu medium plates with an optimal concentration of aureobasidin A (AbA) to examine any protein-DNA interactions. The empty vector containing recombinant GFP was co-transformed as negative control. All primers used are listed in Table S1.

## Results

### Natural variation of iWUE related parameters

In this study, we used a Minicore rice worldwide diversity panel consisting of 200 rice accessions derived from 97 countries (Figure S1A). To determine the dynamic effects of FL on intrinsic water use efficiency (iWUE) operated by stomatal kinetics, we applied a FL treatment with 40 min duration on Minicore rice panel. In this regard, five iWUE related parameters were defined in this study, including iWUE, iWUE_Ir,_ iWUE_Dr_, iWUE_LL_ and iWUE_FL_ during FL treatments (Figure 1A).

**Figure 1.**
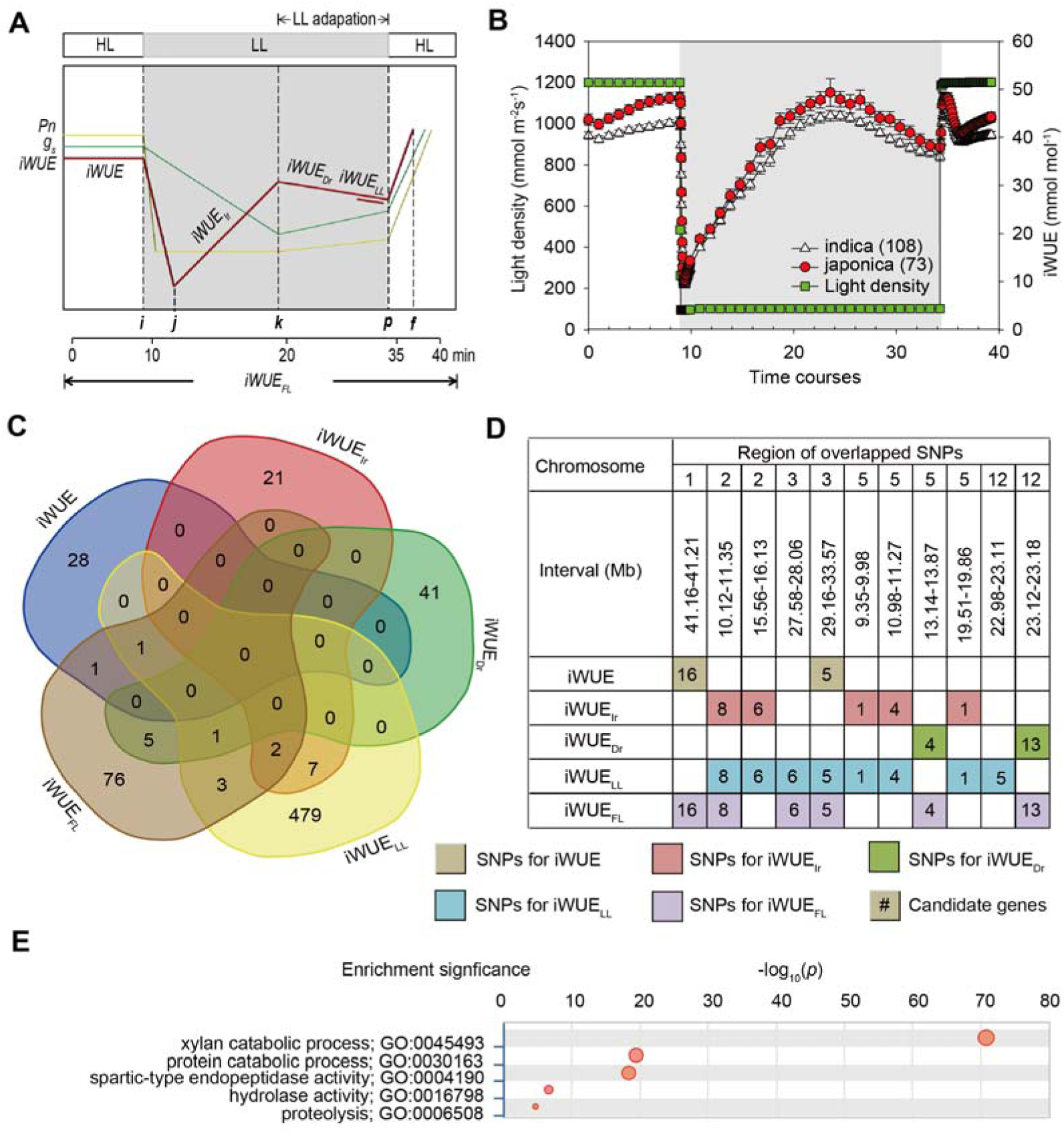
Natural variation of kinetics of iWUE during fluctuating light in Minicore population and candidate genes exploration by genome-wide association study. **A**, A diagram displaying the definition of each intrinsic water use efficiency (iWUE) related traits during fluctuating light (FL) treatment. Parameterization of iWUE relevant traits during the FL treatment. B, Comparison on the dynamics of iWUE between 108 *indica* and 73 *japonica* accessions in Minicore panel during FL treatments. Minicore panel consist of six subpopulations, i.e., ARO, AUS, IND, TEJ, TRJ and admix, while *japonica* include ARO, TEJ and TRJ, and *indica* include IND and AUS. Vertical bars represent mean ± standard error (SE). *n*=108 for *indica* and 73 for *japonica*. **C**, Venn diagram representing the overlapped SNP that significantly associated with five iWUE parameters. **D**, Distribution of overlapped SNPs significantly associated with five iWUE parameters. Different colors represent SNPs significantly associated with each trait. Numbers in cell represents the numbers of candidate genes corresponding to SNPs. **E**, Gene Ontology analysis on the candidate genes across all iWUE parameters.

The values of five iWUE parameters all exhibited normal distribution (Figure S1A-E). In particular, the values of iWUE under high light and low light ranged from 20 μmol mol^-1^ to 70 μmol mol^-1^ in Minicore population. iWUE_Ir_ in average in Minicore population is 5, which is two times higher than iWUE_Dr_. The values of iWUE_FL_ were similar as both iWUE and iWUE_LL_.

Correlation analysis suggests that all five iWUE parameters show strong positive correlation with Pearson correlation coefficient (PCC) > 0.85 between each other in the 206 rice accessions (Figure S2). As expect, iWUE is positively correlated with *A*, while is negatively correlated with *g*_s_ (*p*<0.05). In addition, all five iWUE parameters were negatively correlated with τ_cl_ (half-time of stomatal closure), while positively correlated with both *W*_stomata_ and *L*_stomata_. All five iWUE parameters were positively correlated with δ^13^C. iWUE_FL_ and iWUE_Ir_ were more correlated with *A* compared with other derived iWUE parameters (Figure S2).

### Higher iWUE in *japonica* than that in *indica*

There are great natural variation in five iWUE parameters especially for both iWUE and iWUE_FL_ among different subpopulation (Figure S3A-E). The two iWUE parameters (iWUE and iWUE_FL_) in AUS and IND were relatively lower than that in TEJ and TRJ. This case is true for iWUE_Ir_ as well as iWUE_Dr_. But there is no significant difference in iWUE_LL_ (Figure S3D).

Therefore, we compared the dynamics of iWUE during a 40 min FL treatment in 108 and 73 accessions from *indica* (include AUS and IND) and *japonica* (include TEJ, TRJ and ARO) subgroup in Minicore population (Barnaby *et al*., 2020). The values of iWUE, iWUE_Dr_, iWUE_Ir_, iWUE_LL_ and iWUE_FL_ during FL treatment in TRJ subgroup were 20%∼40% higher than that in IND (Figure 2B; Figure S3A-E). In particular, the values of iWUE_FL_ in TRJ are 43% higher than that in IND (Figure S3E).

**Figure 2.**
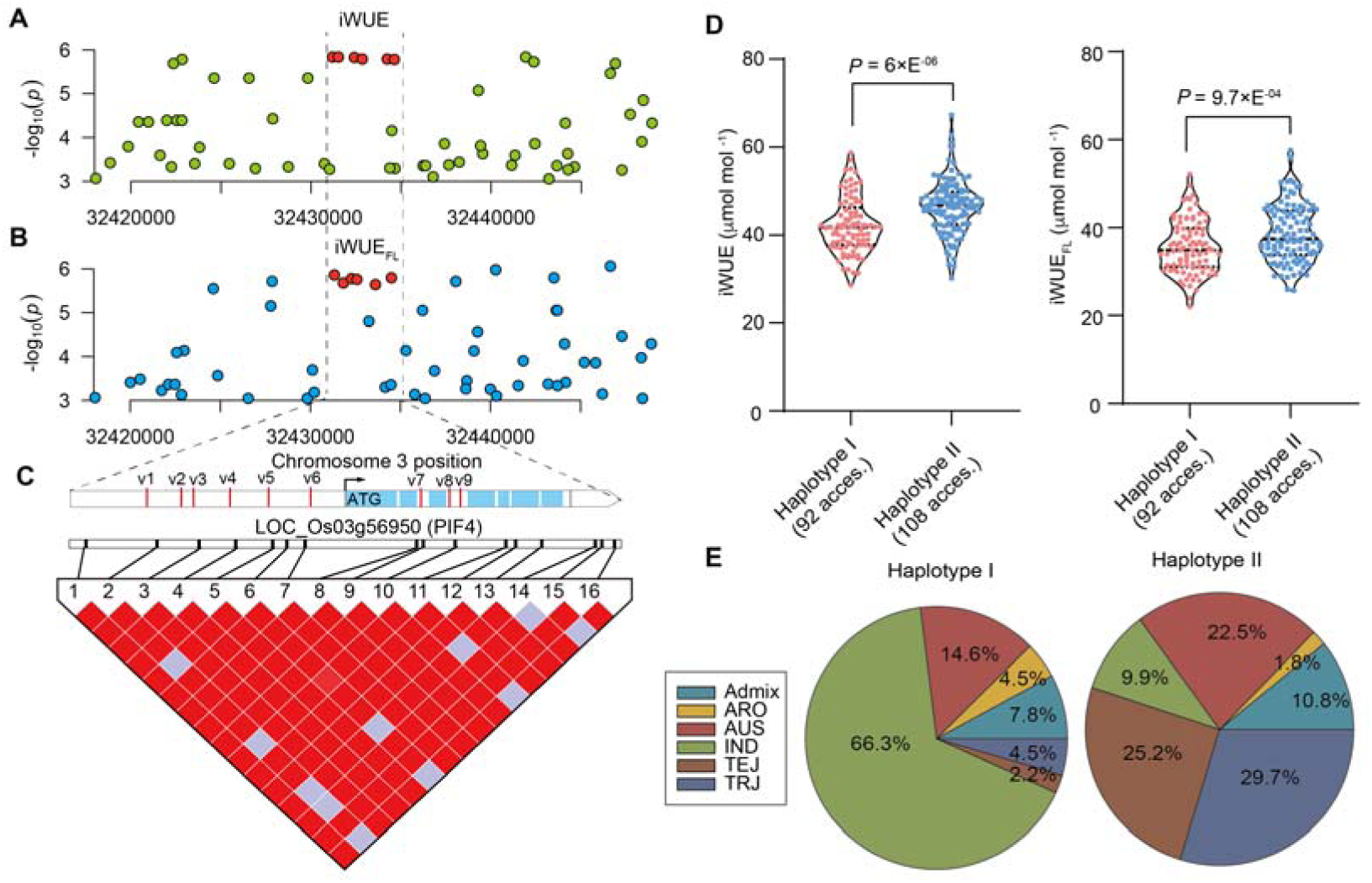
Haplotype analysis of *PIF4* gene. **A-B**, Zoom-out Manhattan plot of iWUE and iWUE_FL_ containing six SNPs in *PIF4* gene. **C**, linkage disequilibrium block analysis of *PIF4* gene. V1-v9 represent the relative position of 9 SNPs on *PIF4* gene. **D**, Distribution of iWUE and iWUE_FL_ in two haplotypes of *PIF4* gene. **E**, Subpopulation distribution in two haplotypes of *PIF4* gene.

To compare the relationship between iWUE_FL_ and drought tolerance, we selected 10 accessions from Minicore population with 5 accession possessing extremely high iWUE_FL_ and 5 accessions possessing extremely low iWUE_FL_. These accessions were exposed to DS for 20 d at seedling stage. Notably, these accessions show similar ranking of iWUE_FL_ that observed in normal condition from previous large-scale iWUE measurements in Minicore population (Figure S4A). Consistently, we found that in these accessions with extreme values of iWUE_FL_, most IND accessions were existed in low iWUE_FL_ group, while most TRJ accessions were in high iWUE_FL_ group (Figure S4B). The growth of these accessions in low iWUE_FL_ group were severely impaired in DS condition. In contrast, plants show relatively tolerance to DS (Figure S4C). The leaf temperature in either normal or DS condition was higher in low iWUE_FL_ accessions than that in high iWUE_FL_ accessions (Figure S4D-E).

### Genome-wide association study on five iWUE parameters

The SNP heritability (*h*^2^_SNP_) of each iWUE parameter was calculated to determine the possibilities of genetic control. Results suggest that *h*^2^_SNP_ in all iWUE parameters were above 0.26 (*P*<0.05). In addition, iWUE_FL_ show highest *h*^2^_SNP_ value (0.51) compared to other four parameters. The value of *h*^2^_SNP_ in iWUE was 0.39 (Table S2). To further determine the candidate genes responsible for the different iWUE parameters, we performed a GWAS using MLM algorithm (Figure S5). Results show that there are two lead SNPs that significantly associated with for three iWUE parameters including iWUE, iWUE_Ir_, and iWUE_Dr_ (Figure S5A-F). There are five lead SNPs that significantly associated with both iWUE_LL_ and iWUE_FL_ (Figure S5G-L). The threshold of significant SNP peaks were determined by 200 times permutation as reported previously (Wang *et al*., 2016). Since strong correlation was observed between iWUE related parameters, we compared the overlapped SNPs that significantly associated with traits.

We found that there are 20 overlapped SNPs in total identified to be significantly associated with different combinations of iWUE related parameters (Figure 1C). These 20 overlapped SNPs were located at Chr. 1, 2, 3, 5 and 12 (Figure 1D). Correspondingly, there are 66 candidate genes that identified in the region of surrounding 50-kb positioned at 20 overlapped SNPs (Table S3). We then performed Gene Ontology analysis, and result show that xylan catabolic process, proteolysis and protein catabolic process in the catalog of biological process, aspartic-type endopeptidase activity and hydrolase activity, acting on glycosyl bonds in the catalog of molecular_function were significantly enriched in the list of 66 candidate genes (Figure 1 E; Table S4).

### Candidate gene responsible for both iWUE and iWUE_FL_

In this study, one lead SNP (3m32427027, *p*_value=4.67E^-06^) on Chromosome 3 was found to be significantly associated with both iWUE and iWUE_FL_. This lead SNP was located in one linkage block window (32.42-32.45 Mb). There are six candidate genes that identified in the window (Figure S6A; Table S3).

We first tested the light response, defined as relative gene expression levels of the six candidate genes under dark / light using 20 rice accessions with 10 extremely high (high iWUE_FL_ subgroup) and 10 extremely low iWUE_FL_ (low iWUE_FL_ subgroup) exposed to DS. Result show that among these six candidate genes, the light responsive expression of only *PIF4* gene was significantly differed between the high iWUE_FL_ group and low iWUE_FL_ group (*p*<0.01) (Figure S6B; Table S5). In addition, the light responsive expression of *PIF4* gene in high iWUE_FL_ group was 40% higher than that in low iWUE_FL_ group (Figure S6B).

### Haplotype analysis and functional validation of *PIF4* gene

According to SNP information in Minicore population, there are 6 SNPs located on the promoter and 3 SNPs located on the intron regions of *PIF4* gene (Figure 2A-B; Table S6). These 9 SNPs belong to one linkage disequilibrium block (Figure 2C). In addition, the 9 SNPs could be divided into two haplotypes, while haplotype I and haplotype II consist of 94 and 112 accessions, respectively (Table S6). The values of both iWUE and iWUE_FL_ in haplotype II were significantly higher than that in haplotype I (Figure 2D). In haplotype I, IND and AUS account 66.3% and 14.6% for total 94 accessions, respectively. In haplotype II, TRJ, TEJ and AUS account 29.7%, 25.2% and 22.5% for 112 accessions, respectively (Figure 4E). We therefore defined hapltype I and haplotype II as Hapl. L and Hapl. H, respectively.

### A casual SNP in PIF4 promoter regulates light response under DS

*PIF4* gene is annotated as phytochrome-interacting factors based on Rice genome annotation database (http://rice.uga.edu/). The PIF4 protein was localized in nuclear as observed by transient transformation in tabacco leaves (Figure 3A). Polygenetic tree analysis suggests that PIF4 (LOC4334324) has less similarity of protein sequenes with other phytochrome-interacting factors genes in rice genome (Figure 3B). In addition, the *PIF4* was highly expressed in dark and shows gradually decreased expression following HL treatment as shown by luciferase signal in rice protoplast (Figure 3B). In particular, the promoter of Hapl. H shows more sensitive to HL than that of Hapl. L (*p*<0.01) (Figure 3B). Furthermore, we determined the activities of promoters harboring 6 SNPs at promoter region of *PIF4* gene in response to light by relative luciferase signal by introducing mutation to Hap. H for different SNPs. Results show that the third SNP (v3) located at -1,075 bp from start codon of *PIF4* gene has greatest sensitivity to HL (Figure 3D).

**Figure 3.**
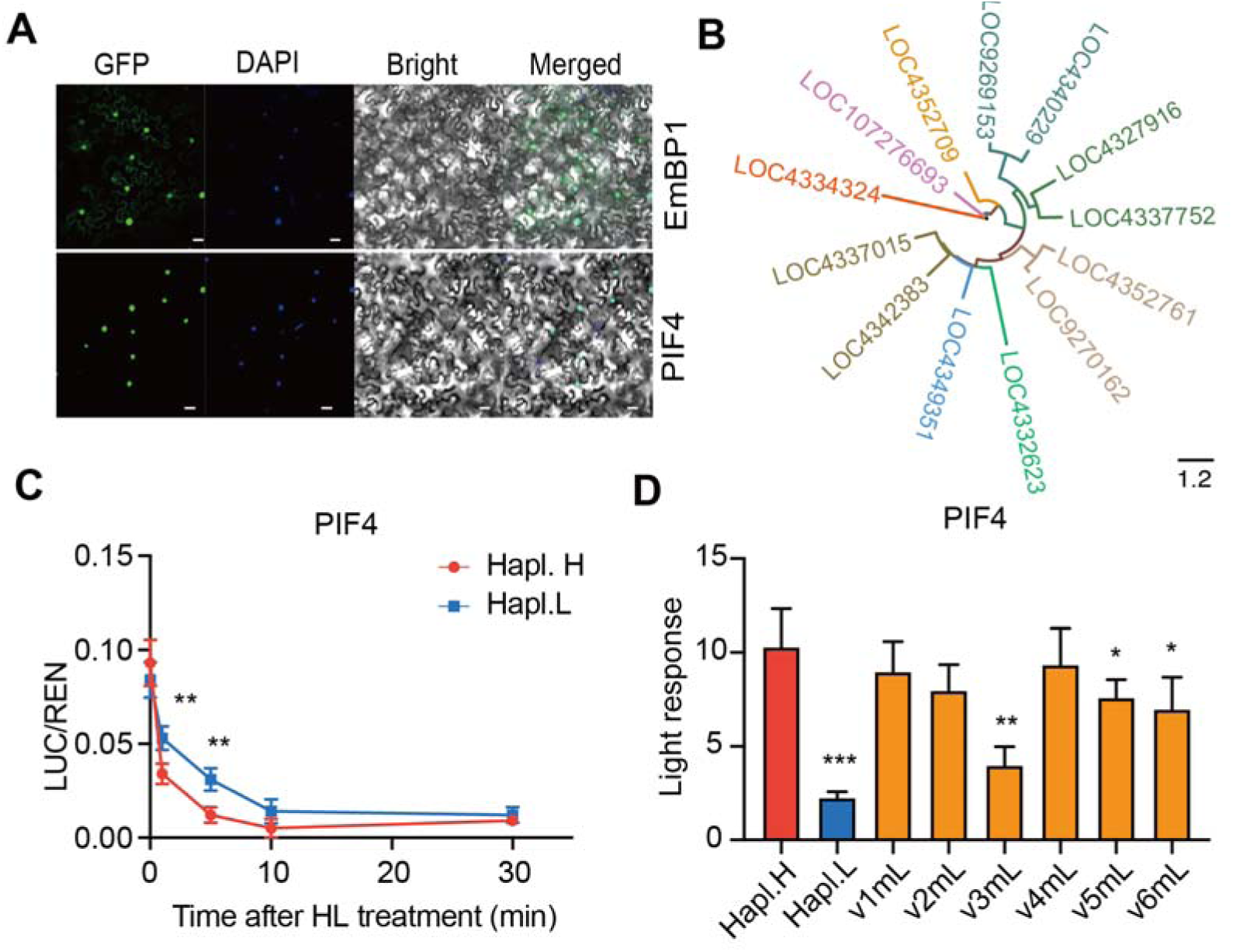
Gene expression and subcelullar localization of PIF4. **A**, OsPIF4 protein is located in nuclear. Subcellular localization was performed using transient transformation in tobacco leaves. A known bZIP transcription factor *EmBP1* gene was used as positive control. **B**, Polygenetic tree of *PIF4* and other 12 Phytochrome-interacting factors genes in rice genome based on protein alignment from NCBI database. **C**, Response of *PIF4* promoter activity to light switch from dark to high light for different duration. Haplotype H and Haplotype L represent the two versions present in *PIF4* promoter. Protoplast were exposed to dark for 24 h, and followed by 30 min high light. **D**, Light response analysis of different variations in *PIF4* promoter. v1mL-v6mL represent six mutation versions introduced to haplotype H of *PIF4*. Data are mean ± s.d. (*n* = 3 biologically independent samples).

The function of *PIF4* gene was further validated using gene-editing technique. A modern rice cultivar WYG7, belong to Hap. H of *PIF4*, were used as background, and the *PIF4* gene was knock-out via CRISPR/CAS9, leading to four nucleotides “GATG” deletion containing v3 SNP (named PIF4^v3m^) (Figure 4A). The growth of PIF4^v3m^ rice line was severely impaired in DS condition, with ∼42% reduction in gene expression in PIF4^v3m^ line (Figure 4B-C). In contrast, we created three overexpression rice lines (OE) with 1.5 times higher expression compared to WYG7 (Figure 4D). Plants of three OE lines in moderate DS were grown better than that of WYG7 (Figure 4E). Both of iWUE and iWUE_FL_ show at least 20% reduction in PIF4^v3m^ compared to WYG7 (Figure 4F-G). The two parameters were stimulated at least 25% in three OE rice lines compared to WYG7 (Figure 4H; Figure S7A). In addition, some photosynthetic traits and agronomic traits were increased by ranging from 10% to 57% in OE rice lines, including chlorophyll contents, photosynthesis rates (Pn), tillering and yield (Figure S7B-F). Consistently, we found that these photosynthetic traits and agronomic traits were declined by ranging from 9% to 48%, while τ_cl_ representing half-time of stomatal closure was increased 20% in PIF4^v3m^ relative to WYG7 (Figure S7F-J).

**Figure 4.**
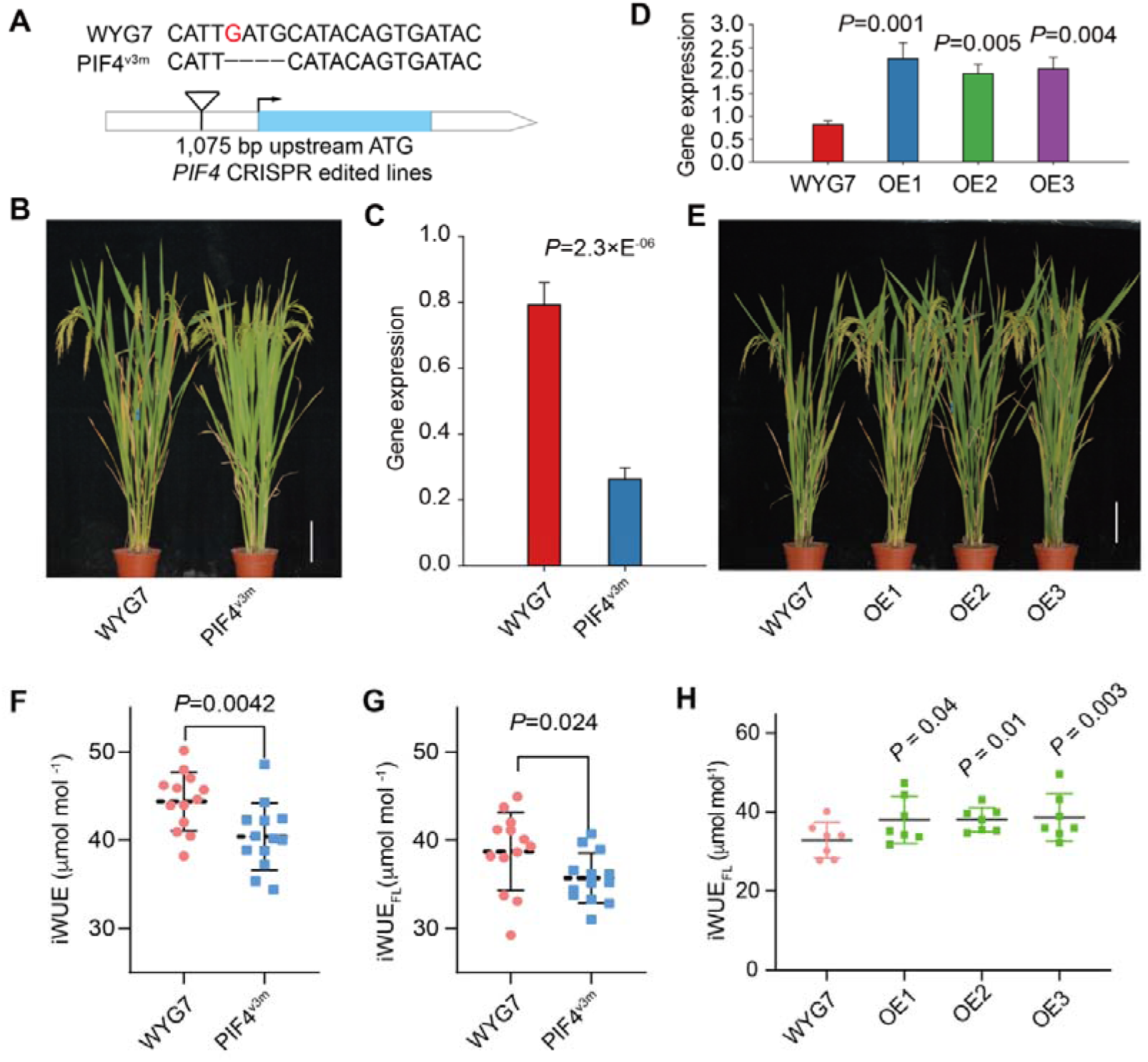
*PIF4* regulates iWUE and iWUE_FL_ in drought stress condition. **A**, Four nucleotides “GATG” including v3 SNP were deleted by CRISPR/CAS9 in PIF4 line (named PIF4^v3m^). **B**, Imaging of WYG7 and PIF4^v3m^ in drought stress (DS) condition. Plants grown in the paddy field, and moved out for imaging at grain filling stage. White vertical bar represents 10cm. **C**, Gene expression levels of *PIF4* in WYG7 and PIF4^v3m^ line. **D**, Gene expression levels of *PIF4* in WYG7 and *PIF4* overexpression (OE) lines. **E**, Imaging of WYG7 and three *PIF4*-OE lines. Plants grown in the paddy field, and moved out for imaging at grain filling stage. White vertical bar represents 10cm. **F-G**, iWUE and iWUE_FL_ in WYG7 and PIF4^v3m^ rice lines in DS condition. **H**, iWUE_FL_ in WYG7 and three *PIF4*-OE rice lines in DS condition.

### PIF4 is involved in SAL1-PAP retrograde signaling pathway under DS

To further elucidate the molecular mechanism of PIF4 regulating iWUE and drought tolerance, we performed a combined analysis of transcriptome and metabolites. Results show that the amounts of 130 metabolites were differentially altered, including 66 downregulated and 64 upregulated metabolites in PIF4^v3m^ rice relative to WT exposed to DS (Table S7). In particular, adenosine 3’,5’-diphosphate (PAP) was decreased by 85% in PIF4^v3m^ rice. The results were further confirmed by targeted metabolites and increased by 94% in two OE lines of *PIF4* gene (Figure 5A). SAL1-PAP retrograde pathway is known to interact with ABA signaling to regulate stomatal closure and drought resistance (Pornsiriwong *et al*., 2017). In consistent with this, we observed a dramatical increase in PIF4^v3m^ rice line based on transcriptome analysis (Figure 5B). The gene expression of *SAL1* is negatively associated with *PIF4* gene expression in PIF4^v3m^ and OE rice lines (Figure 5C). In addition, we found that PIF4 directly binds to G-box motif of *SAL1* at -389 bp (Figure 5D). Luciferease assays and yeast-one hybrid assays suggest that PIF4 is translationally inhibit the expression of *SAL1* gene (Figure 5E; Figure S8A-B).

**Figure 5.**
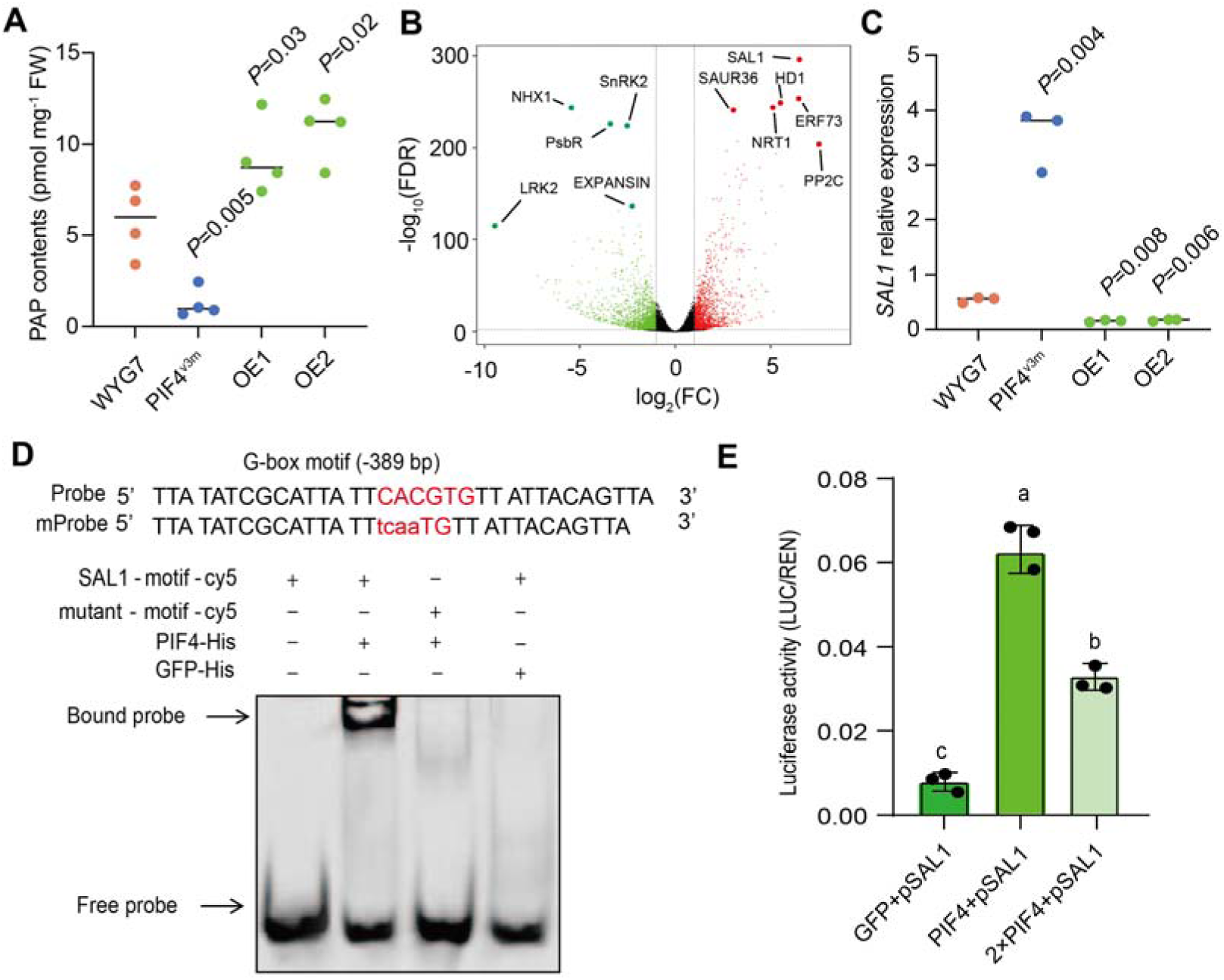
PAP contents and differentially expressed genes analysis in WYG7 and *PIF4* transgenic rice lines exposed to drought stress in the field. **A,** PAP contents in WYG7 and PIF4 transgenic rice lines (PIF4^v3m^ and *PIF4*-OE). **B**, Volcano plot representing differentially expressed genes (DEGs) between WYG7 and PIF4^v3m^. **C**, *SAL1* gene expression via qPCR against *actin*. **D**, EMSA for PIF4 binding to G-box motif of *SAL1* promoter. **E**, Luciferase activity determination for PIF4 transcriptionally repress the expression of *SAL1*. *n*=3 for panels **C, E**.

We hence constructed co-expression of *PIF4*-*SAL1* rice lines, and results show that OE lines of *SAL1* (*SAL1*-OE) was severely impaired under DS condition, while the protein levels of SAL1 were increased more than 10-time in *SAL1*-OE line relative to WYG7 (Figure 6A-B). In contrast, the growth in co-expression line of *PIF4* and *SAL1* (*PIF4*-*SAL1*) were dramatically better than that in WYG7 and *SAL1*-OE, but less pronounced than *PIF4*-OE, which could mainly due to inhibitory effects of *SAL1* under DS condition (Figure 6A-B). The PAP contents were dramatically increased in *PIF4*-OE rice line, while decreased in *SAL1*-OE line (Figure 6C). The WUE_FL_ in *PIF4*-*SAL1* line was 15% decreased relative to *PIF4*-OE, but increased ∼26% compared to WYG7 (Figure 6D), which leads to similar ranking of drought tolerance as shown by tillering and yield (Figure 6E-F).

**Figure 6.**
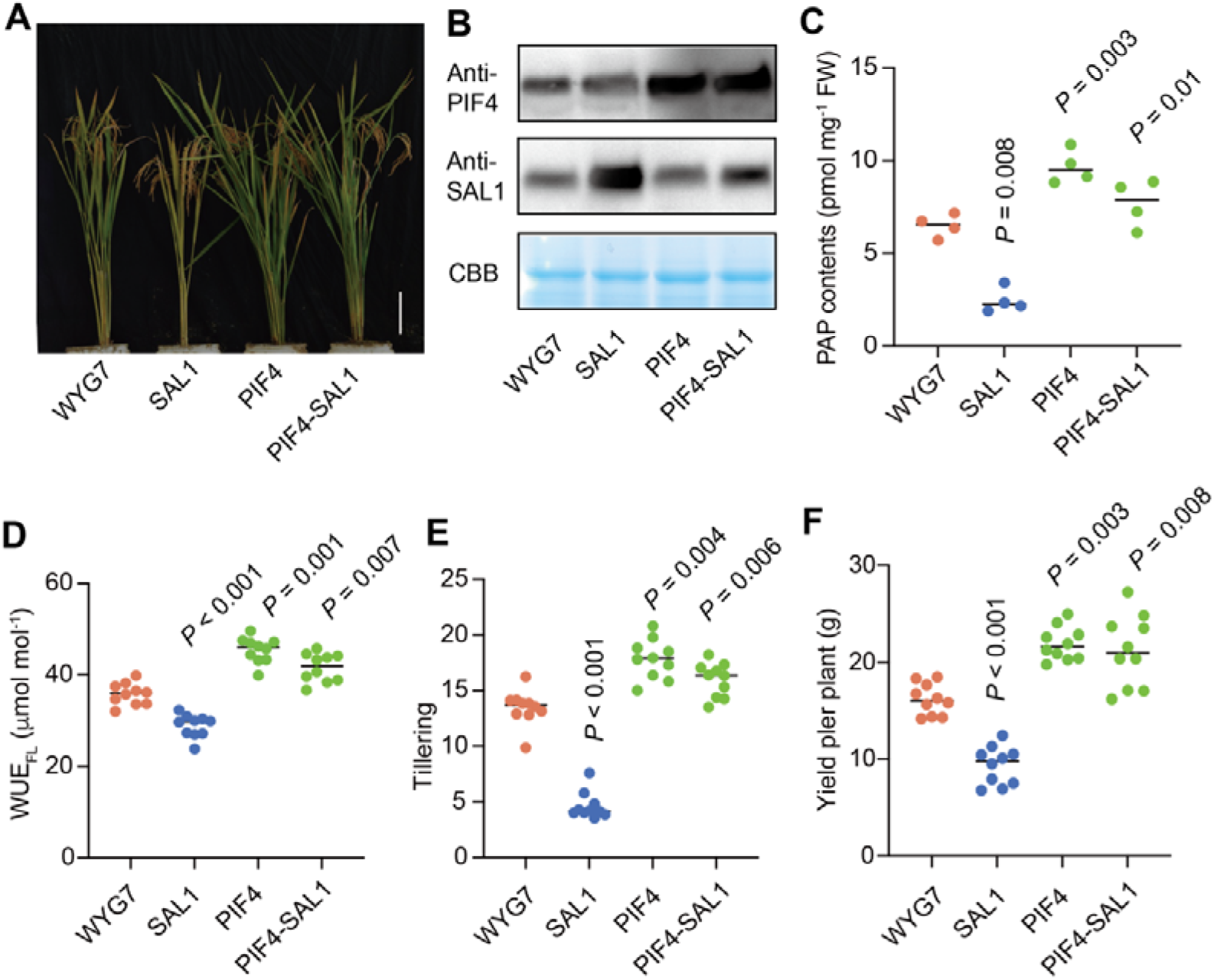
PIF4 compensates the drought sensitivity caused by SAL1 in the field. **A**, Images of WYG7, overexpression lines of SAL1 (*SAL1*-OE), PIF4 (*PIF4*-OE) and co-overexpression of both *SAL1* and *PIF4* (*PIF4*-*SAL1*) exposed to DS at graining stage. **B**, Protein expression levels in WYG7 and different overexpression lines of *SAL1* and *PIF4*. **C-F**, PAP contents, iWUE_FL_, tillering and yield in WYG7 and different overexpression lines of *SAL1* and *PIF4*. Each bar data represents the mean of replicates (*n*=10) ±SE for panel **C-F**. One-way *ANOVA* was used to determine the significance level between WYG7 and these overexpression lines of *SAL1* and *PIF4*.

### PIF4 regulates stomatal closure speed by transcriptional repression to NHX1

NHX1 is a potential regulator that responsible for stomatal dynamics during DS, and based on transcriptome, we found that the expression of *NHX1* is downregulated by 85% in PIF4^v3m^ under DS. EMSA experiments show that PIF4 could transcriptionally activate the expression of *NHX1* through direct binding to G-box motif of *NHX1* at ∼392 bp from start codon (Figure S9A-B). Y1H assays also show that PIF4 specifically binds to G-box of motif present in promoter of *NHX1* (Figure S9C). Furthermore, we constructed the co-expression lines of *PIF4* and *NHX1* (*PIF4*-*NHX1*). Results show that *PIF4*-*NHX1* performs stay-green phenomenon but not WYG7 grown in DS in the field, as also shown 25% increase in chlorophyll contents in *PIF4*-*NHX1* (Figure 7A-B), which ensures better photosynthetic capacity and iWUE kinetic parameters during FL (Figure 7C-D; Figure S10-A-B). The stomatal close speed was also enhanced in *PIF1*-*NHX1* as shown lower values of τ_cl_ compared to WYG7 during FL (Figure S10C). The changes of these parameters lead to increased drought tolerance as shown by higher tillering and yield in *PIF4*-*NHX1* than WYG7 exposed to DS (Figure 7E-F). There is no significant differences between *PIF4*-*NHX1* and *PIF4*-OE in WUE_FL_ and drought tolerance (Figure S10D-E).

**Figure 7.**
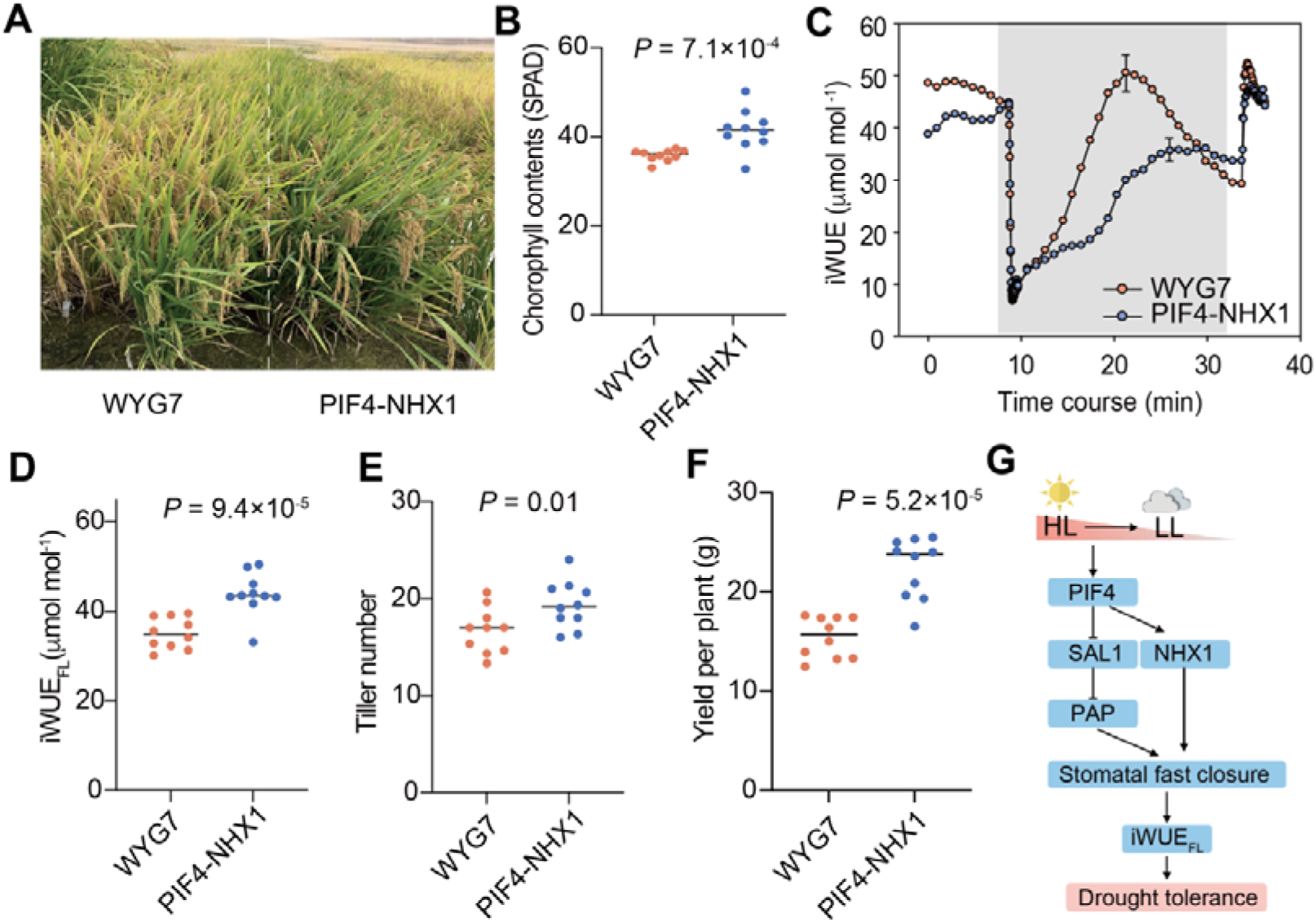
PIF4 promotes *NHX1* gene expression enhancing drought tolerance. **A**, Field images of WYG7 and co-expression of *PIF4* and *NHX1* (*PIF4*-*NHX1*) at graining stage. **B**, Chlorophyll contents between WYG7 and *PIF4*-*NHX1*. **C**, Dynamics of iWUE during FL between WYG7 and *PIF4*-*NHX1*. Vertical bar represent maximum standard errors. **D-F**, iWUE_FL_, tiller number and yield per plant between WYG7 and *PIF4*-*NHX1*. Each bar data represents the mean of replicates (*n*=10) ±SE for panel **B-F**. One-way *ANOVA* was used to determine the significance level between WYG7 and *PIF4*-*NHX1*. **G**, Summarized model representing the regulation of PIF4 on iWUE during fluctuating light improving drought tolerance via cooperating with *SAL1* and *NHX1*.

Collectively, light when switched from HL to LL, *PIF4* gene expression was stimulated, which leads to increased *NHX1* gene hence fast stomatal closure, simultaneously, PIF4 transcriptionally repress the expression of *SAL1*, resulting in PAP accumulation and stomatal fast closure. The two pathways regulated by *PIF4* (*PIF4*-*SAL1* and *PIF4*-*NHX1*) eventually promotes iWUE during FL, and drought tolerance (Figure 7G).

## Discussion

Plants always exposed to a naturally changing light environment due to many factors, e.g., occasional cloud-cover, called sunflecks or shadeflecks (Lawson & Blatt, 2014). iWUE is mainly controlled by *A* and *g*_s_, which causes the complex changes of iWUE during FL. We found that the values of iWUE were gradually increased started from 1 min after the switch from high to low light until 15 min. During this period, *g*_s_ decreased continuously and *A* was unchanged (Qu *et al*., 2016). Therefore, the iWUE_Ir_ is mainly driven by τ_cl_ (time constant of stomatal closure speed) as documented previously (Qu *et al*., 2020), which leads to high iWUE_FL_ accompanied with fast stomatal closure in rice population (Figures S2).

Recently, still limited numbers of genes regulating photosynthetic efficiency under high light were identified, such as NRP1 (Chen *et al*., 2021) and *EmBP1* (Perveen *et al*., 2020), as well as under FL, including *PsbS* (Kromdijk *et al*., 2016). However, the genes related to dynamics of iWUE under FL were less reported. Based on GWAS, we found that 20 SNPs were overlapped across different iWUE parameters. Consistently, we found that the candidate gene *RM5521* at chromosome 2 (Adachi *et al*., 2019) was also observed in the linkage disequilibrium block (15.56-16.13 Mb) that associated with both iWUE_Ir_ and iWUE_LL_ (Figure 3B). In addition, we found that *OsCML4* (29.16-30.57Mb) at chromosome 3 (Yin *et al*., 2015) was also identified in the list of candidate genes responsible for iWUE, iWUE_LL_ and iWUE_FL_ (Figure S5A, G, I). *OsSLAC1* at chromosome 4 (Yamori *et al*., 2020) was identified in the iWUE_LL_ (Figure S5G). These suggest that the genetic control of iWUE during FL is closely related to stomatal features.

SNP heritability (*h*^2^_SNP_) is an important parameter to determine the proportion of genetic control of some complex traits. It was reported that *h*^2^_SNP_ of most ecophysiological traits, such as *A*, *g*_s_, iWUE, dark respiration (*R*_d_) are 0.01∼0.6, while *h*^2^_SNP_ of morphological traits include biomass, tillering and plant height were around 0.5∼0.8 (Qu *et al*., 2017). In our study, we found that iWUE_FL_ has highest *h*^2^_SNP_ (0.5) among the five iWUE parameters (Table S2). Based on GWAS, we found that the allelic variation of PIF4 is strongly associated with iWUE_FL_ and drought tolerance. PIF4 belong to phytochrome-interacting factors gene family, which is known to mediate the photomorphogenic response in *Arabidopsis thaliana* (Lau & Deng, 2010).

PIFs are a small subfamily of basic helix-loop-helix (bHLH) transcription factors that play multiple functions in processes such as shade avoidance responses and temperature through regulating chlorophyll pigments biosynthesis and hypocotyl elongation (Leivar & Monte, 2014; Ma *et al*., 2016). In this study, we found the mutant of *PIF4* (PIF4^v3m^) shows lower chlorophyll contents than WYG7 during FL, suggesting the complex regulation of chlorophyll biosynthesis under FL coincident with DS, which might be due to compensated expression from homologs in other gene family members (Qu *et al*., 2020). We also found dramatic decline in gene expression in PIF4^v3m^, revealing that the position of SNP (v3) is very likely to be regulated by some transcription factors. PIFs accumulate in the dark to promote skotomorphogenesis, whereas light induces the rapid phosphorylation and degradation of PIFs, through the activity of phyB and phyA in Arabidopsis (de Lucas *et al*., 2008; Jang *et al*., 2010). Until now, there has been little information on PIFs in crops. In our study, we found that the sequence of PIF4 protein was distinct from other PIFs homologs in rice, and it shows decreased gene expression followed by HL exposure from dark (Figure 3B-C), suggesting like PIF1, activated phyB interacts with PIF4 and induces its phosphorylation and degradation under light irradiation (Xu & Zhu, 2021).

Previous molecular studies showed that PIFs directly regulate the expression of downstream genes by binding to a G-box motif (CACGTG) present in their promoters, including ABI5 and CDF2 (Kim *et al*., 2016; Gao *et al*., 2022). In our study, we found PIP4 directly binds to G-box motif of both *SAL1* and *NHX1* with differentially transcriptional regulation. PAP is thought to act as a typical retrograde signal, which is produced in chloroplasts under drought stress to induce the expression of nuclear-encoded stress response genes, leading to stomatal closure (Pornsiriwong *et al*., 2017). Based on combined analysis of transcriptome and metabolism, we found that PIP4 promotes biosynthesis of PAP, through inhibiting *SAL1* expression under DS, suggesting that PIP4 is probably involved in SAL1-PAP mediating chloroplast retrograde signaling pathway. Therefore, we found that PIP4 regulated this pathway could also participate in stomatal driven physiological response to FL and DS in rice, in addition to known function that altered nuclear expression during HL and DS in Arabidospsis (Estavillo *et al*., 2011).

Stomatal movements rely on alterations in guard cell turgor. We found that another gene underlying stomatal closure, *NHX1* during FL as observed in previous study (Qu *et al*., 2020), could also be promoted by PIF4 by binding *NHX1* G-box motif. *NHX1* is a Proton Sodium exchanger, overexpressing *NHX1* is proved to improve the drought and salt tolerance in Arabidopsis (Brini *et al*., 2007) as well as in other species. In our study, we found co-overexpression of *PIF4* and *NHX1* could dramatically stimulate plant growth under DS through accelerating stomatal resonse speed during FL, suggesting *PIP4*-*NHX1* modules could contributes to accumulation of K^+^ into the vacuole of plant cells, thereby increasing their osmotic potential and driving the uptake of water that generates the turgor pressure necessary for cell expansion and growth under DS (Barragan *et al*., 2012).

## Conclusion

iWUE plays important roles in drought tolerance especially under FL. In this study, we investigated the natural variation of five iWUE kinetic parameters during FL in 200 rice accessions. Results show that iWUE_FL_ has closely related with drought tolerance with moderately high SNP heritability. GWAS identifies six candidate genes, a *PIF4* gene show significantly greater sensitivities to light in iWUE_FL_ subgroup. A SNP (v3) at promoter region of *PIF4* mainly contributes to its declined expression under light. Knocking out v3 SNP results ∼20% reduction in iWUE_FL_. PIF4 transcriptionally represses and activates *SAL1* and *NHX1*, respectively, through binding to G-box motifs of the two genes. We proposed that PIF4 promotes iWUE_FL_ through stomatal fast closure resulted by decreased *SAL1* and increased *NHX1* gene expression during FL, eventually facilitating to drought resistance.

## Data and materials availability

All data is available in the manuscript or the Supporting Information. Thus, this document includes 10 Figures and 8 Tables as supplementary materials.

## Acknowledgements

We thank Prof. Chengcai Chu of South China Agricultural University for sharing Minicore seeds. This work was supported by National Natural Science Foundation of China (32170245; 32260447), Natural Science Foundation of Zhejiang Province (LQ20C130003), Sanya Yazhou Bay Science and Technology City (SCKJ-JYRC-2022-04), Scientific Research Fund of Zhejiang Provincial Education Department (YZ0Z145972), Huzhou public welfare application research project (2021GZ26) and National Training Programs of Innovation and Entrepreneurship for Undergraduates (2022hzxy019).

## Author contributions

Conceptualization, M.Q., Z.X.; Methodology, S.L., J.E., Y.L.; Investigation, S.L., Y.L., C.L., F.Z.; Writing, M.Q., Z.X., J.E.; Funding Acquisition, M.Q., S.L.; Resources, M.Q.; Supervision, M.Q.

