## Supplementary Figures for "PIF4 promotes water use efficiency during fluctuating light and drought resistance in rice"

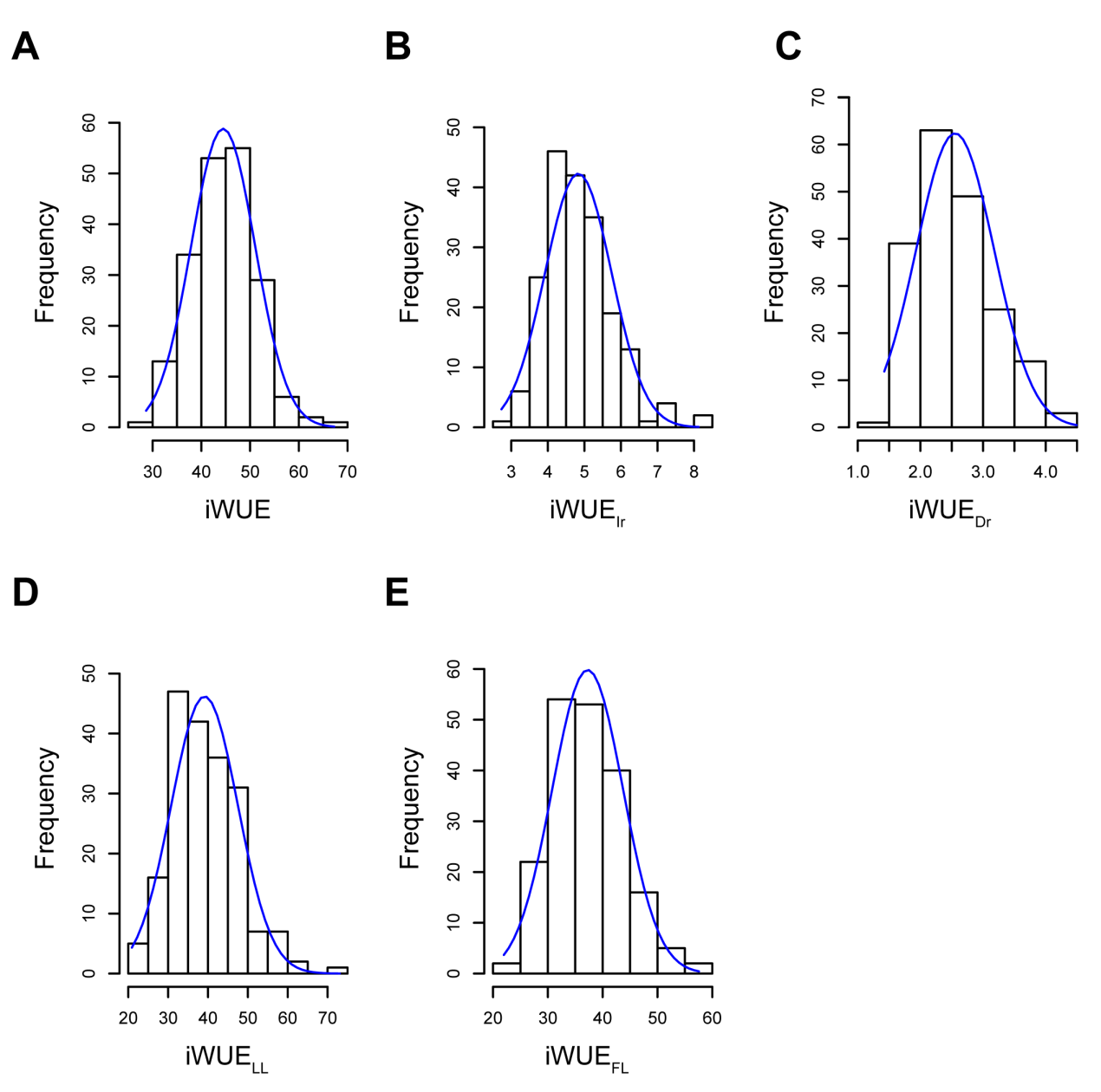

**Figure S1.** Distribution of five iWUE parameters in the rice Minicore population consisting of 200 accessions. **A-E**, iWUE, iWUE_Ir_, iWUE_Dr_, iWUE_LL_, and iWUE_FL_.

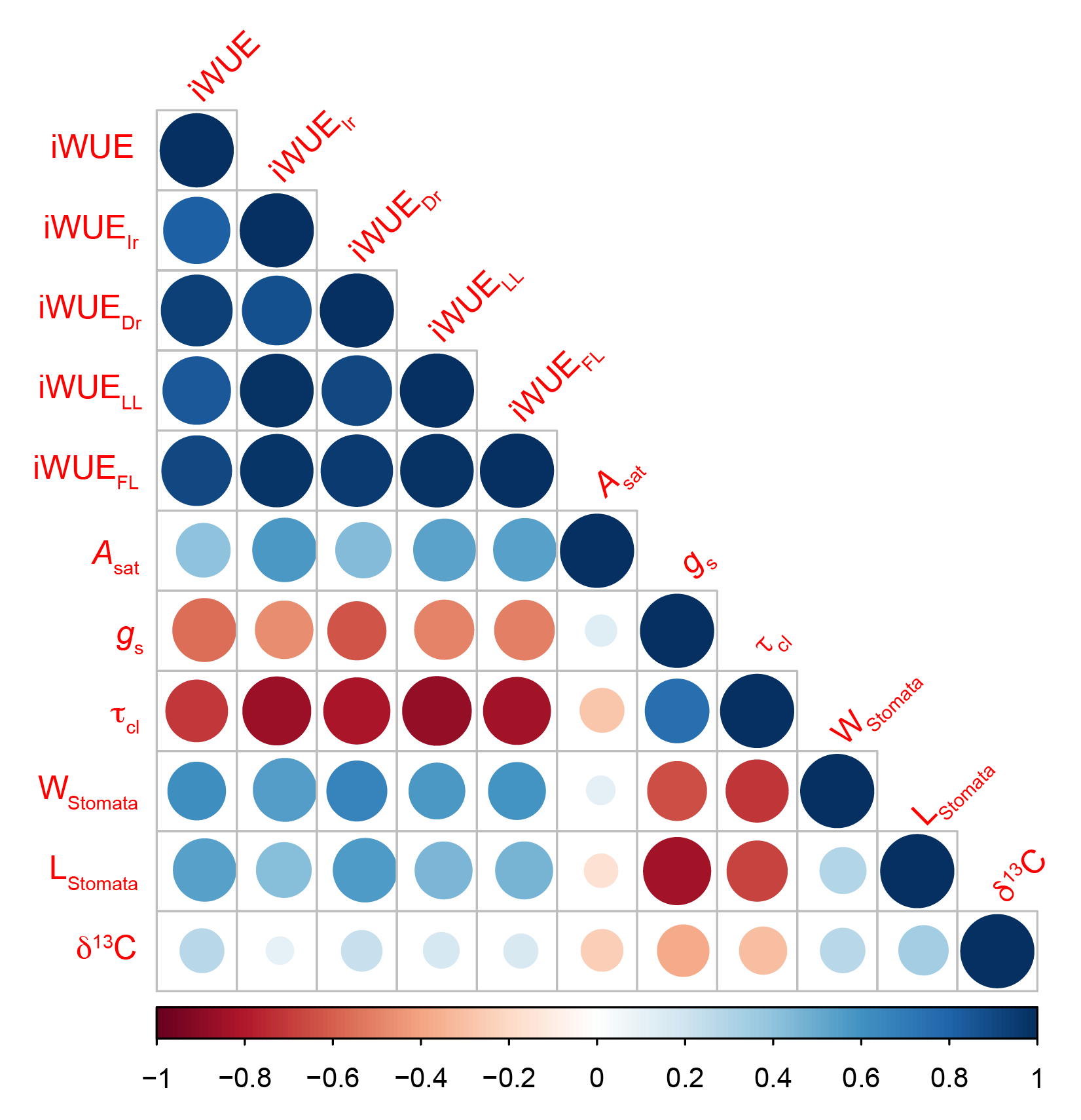

**Figure S2.** Association analysis on five iWUE parameters and other physiology traits in Minicore population under drought stress conditions. The color from blue to red represent the Pearson's correlation coefficient (PCC) ranges from 1 to -1.

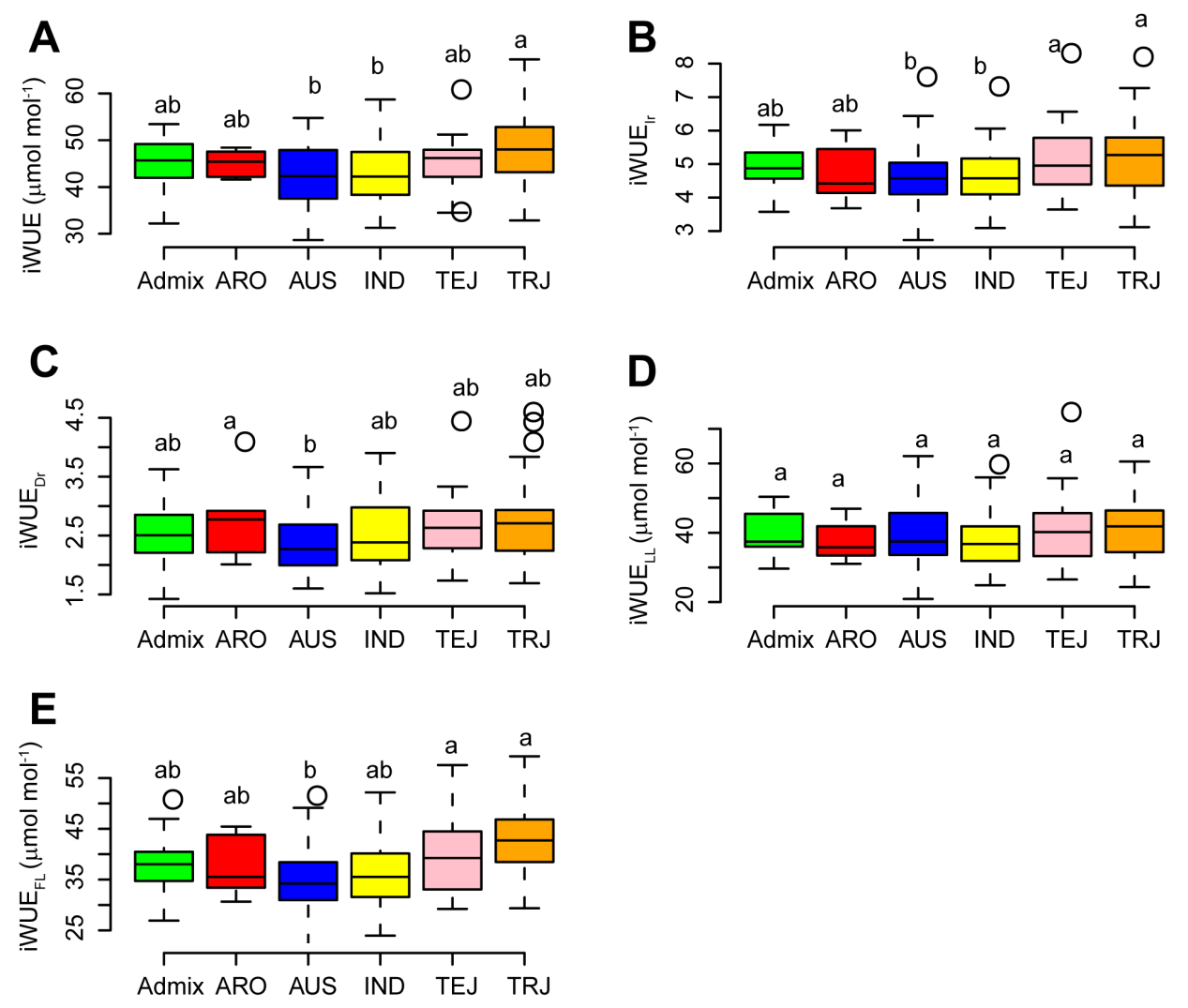

**Figure S3.** Distribution of five iWUE parameters in the Minicore population. **A-E**, iWUE. iWUE_Ir_. iWUE_Dr_. iWUE_LL_ and iWUE_FL_, respectively. In boxplot, the box edges represent the upper and lower quantile with median value shown as bold line in the middle of the box. The individual outside of range of the whiskers was shown as open dots. The Minicore population encompasses 19, 6, 38, 70, 30 and 37 accessions for subpopulation of Admix, ARO, AUS, IND, TEJ and TRJ, respectively.

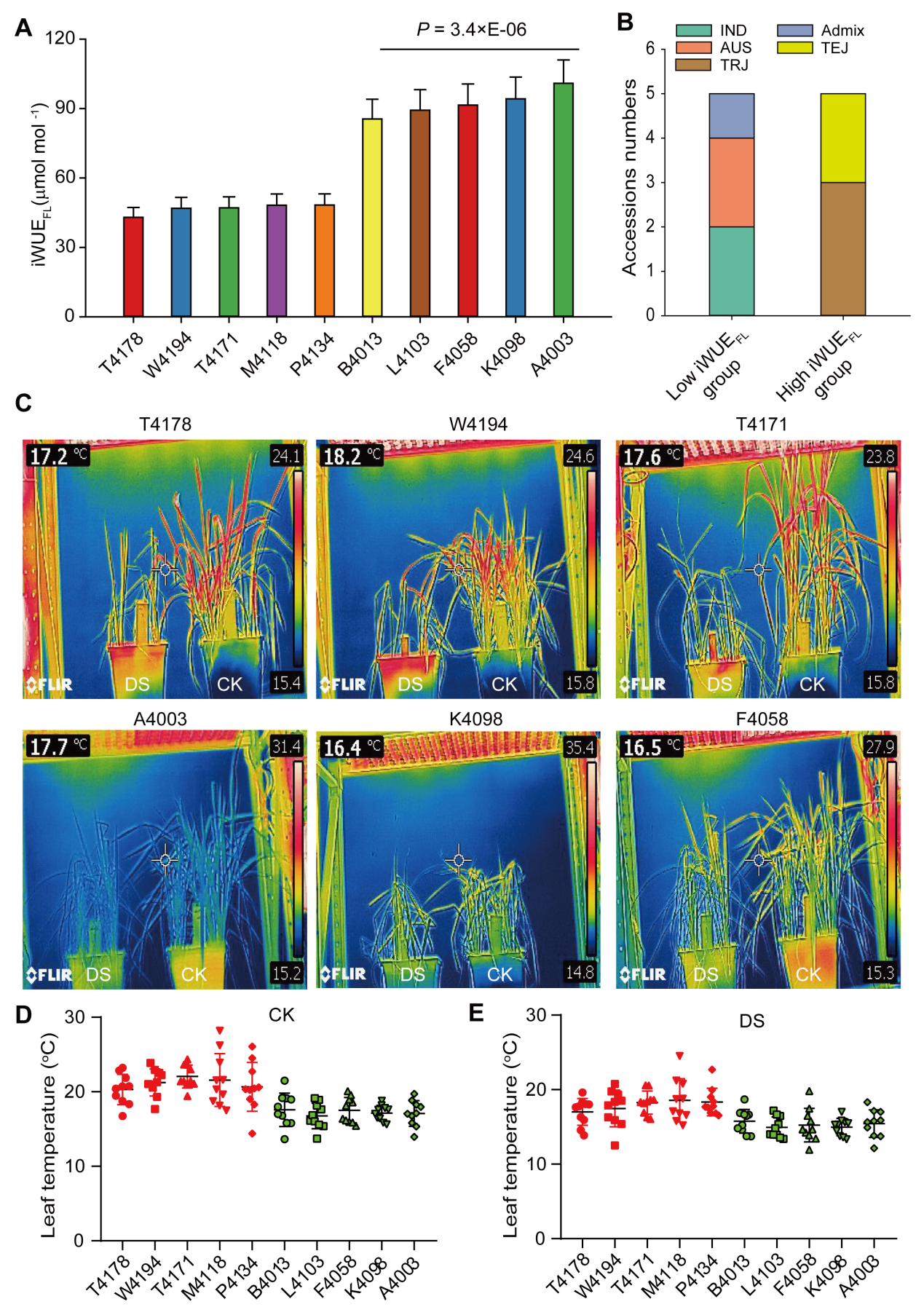

**Figure S4.** iWUE_FL_ is closely related to drought resistance under fluctuating light in rice population. **A**, Comparison on iWUE_FL_ in 10 rice accessions from moderate drought stress (DS) condition. **B**, Accession numbers of each subpopulation in low iWUE_FL_ group and high iWUE_FL_ group. **C**, Thermal imaging of rice plants grown in growth chamber with fluctuating light together with moderate drought stress condition. **D-E**, Canopy leaf temperature in normal (CK) and DS in the 10 rice accessions. Each point represents pixel point extracted from thermal imaging for each rice line.

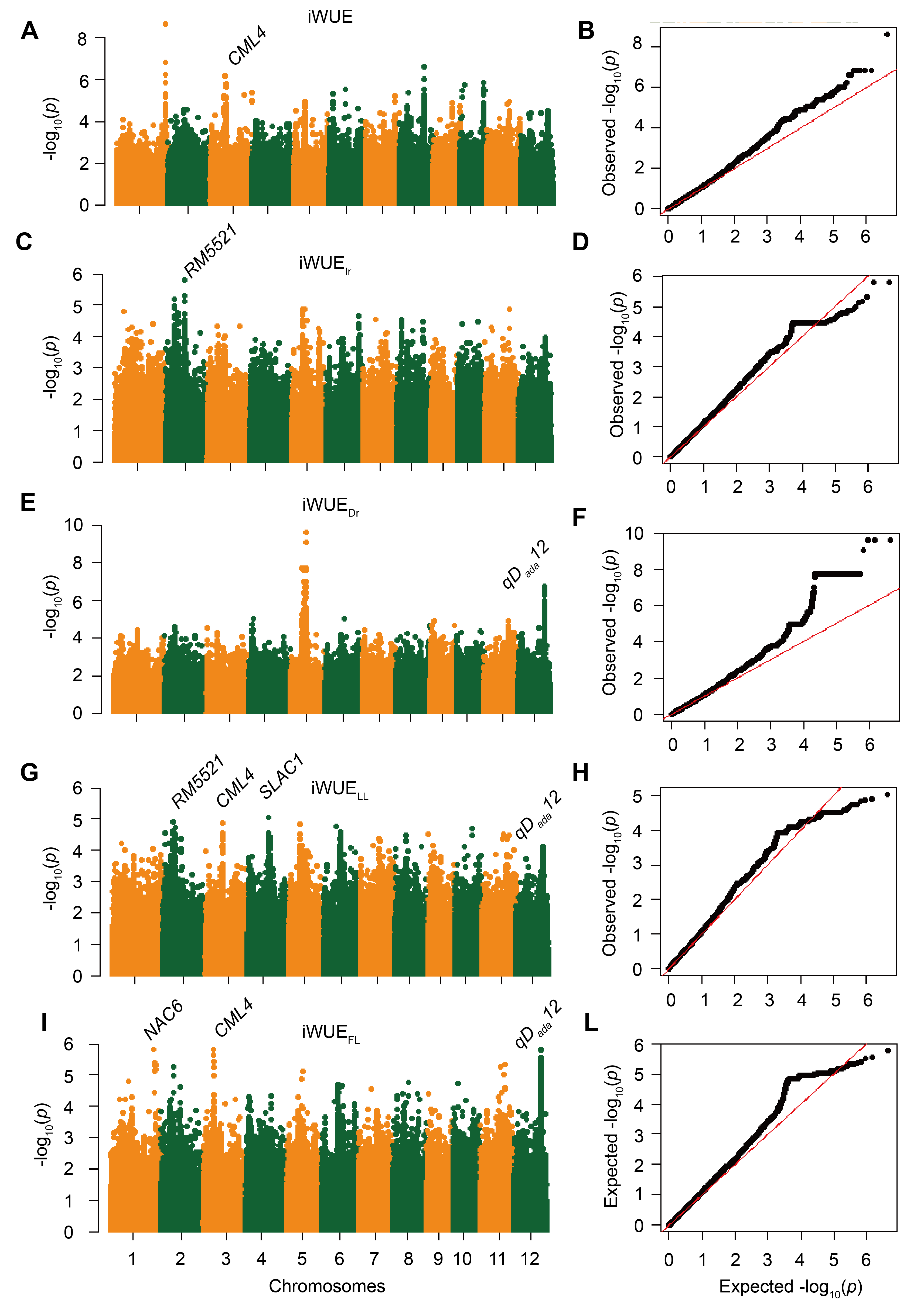

**Figure S5.** Manhattan and QQ plots for different iWUE traits. **A-B**, iWUE; **C-D,** iWUE_Ir_; **E-F,** iWUE_Dr_; **G-H,** iWUE_LL_; **I-L**, iWUE_FL_. Manhattan plots from the association mapping of the five iWUE traits using a linear mixed model (MLM). The QQ plots of the expected versus observed *P-*values of the five iWUE derivated traits. The reported QTLs-based genes were labeled in each panel as well documented earlier including *CML4* (Yin et al., 2015), *RM5521* (Adachi et al., 2019), *NAC6* (Nakashima et al., 2007), *SLAC1* (Kusumi et al., 2012) and *qD_ada_12* (Chen et al., 2020).

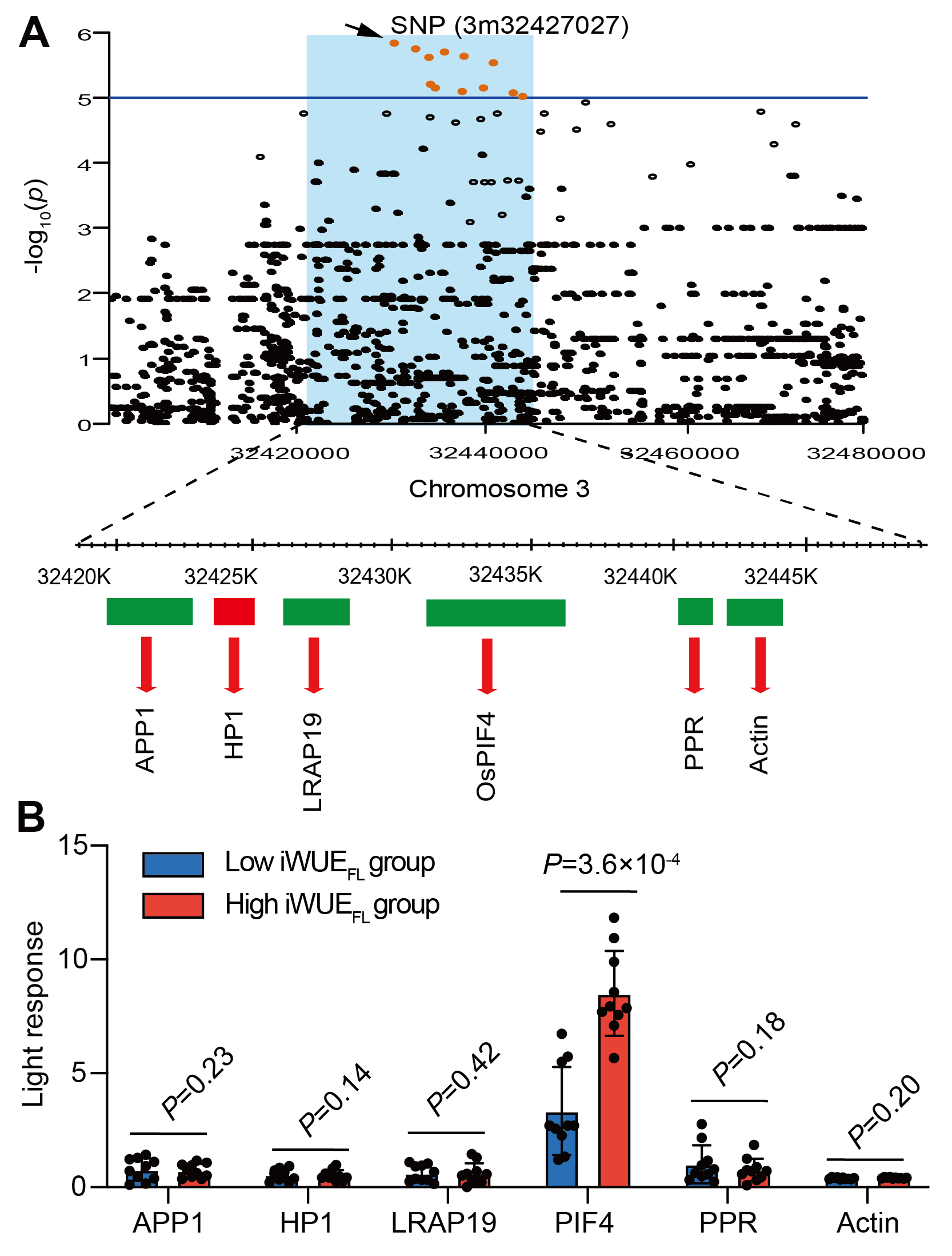

**Figure S6.** Zoom-out plot of Manhattan plot of iWUE_FL_ and differential gene expression analysis of candidate genes. **A**, Zoom-out plot of Manhattan plot of iWUE_FL_ highlighting the lead SNP (3m32427027) and its associated candidate genes surrounding the linkage disequilibrium (LD) block (30.41-41.21Mb). The horizontal line at the P-value 10^-5^ represents the cut-off values for significant SNPs. The detailed information of the candidate genes is encompassed in Table S3. **B**, Light response for candidate genes within the LD block surrounding the lead SNP within two groups possessing contrasting iWUE_FL_. The light response values were determined by expression levels under dark against that under light for each candidate gene. Each group has 10 rice accessions in the Minicore population. *n*=10. A student *t*-test was used to determine the significance level. The expression values of each candidate gene were referred to Table S5.

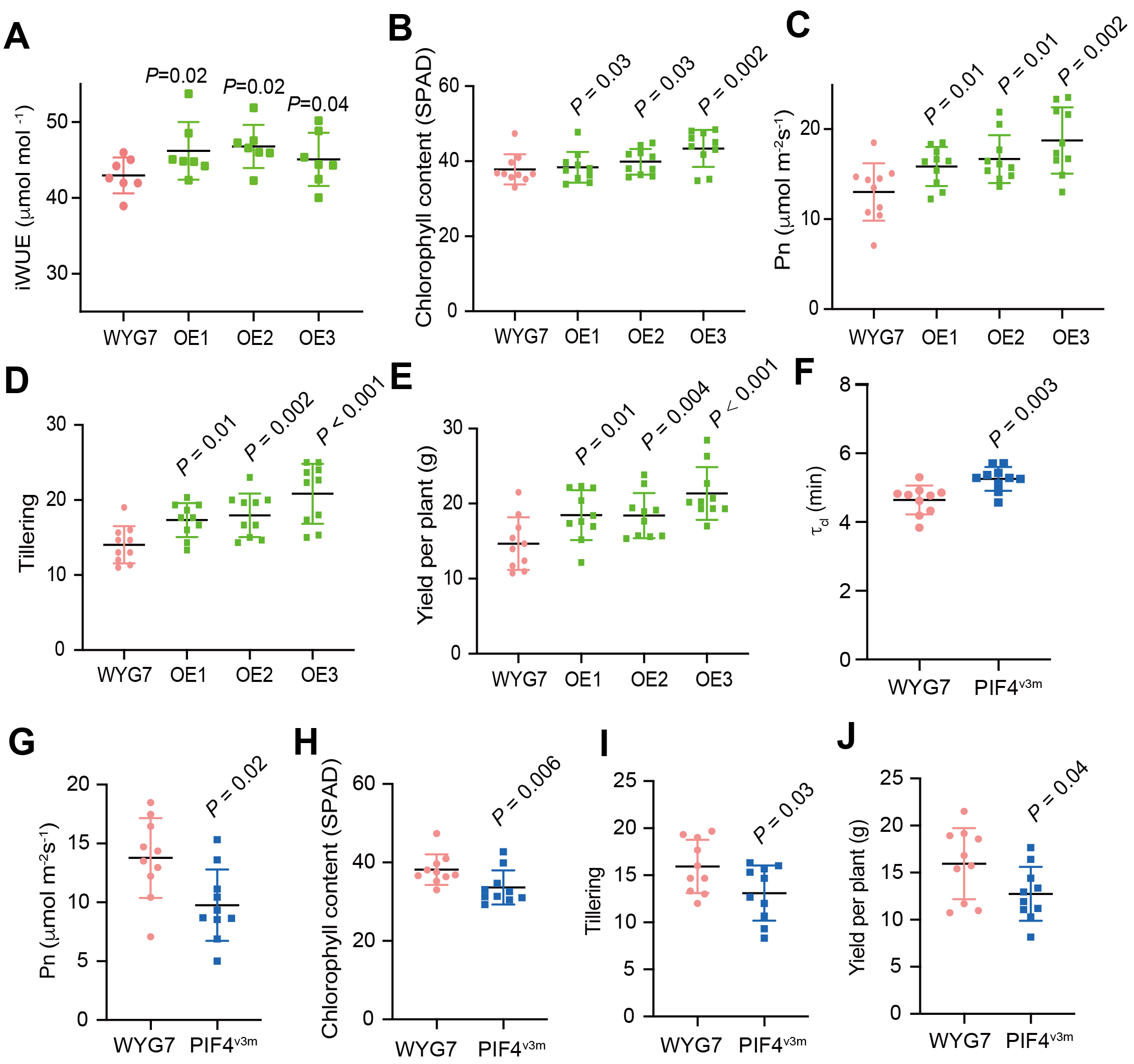

**Figure S7.** Physiological and agronomic traits of *PIF4* overexpression and its CRISPR-edited rice lines. **A-E,** iWUE, chlorophyll contents, Pn, tillering and yield per plant, in WYG7 and PIF4 overexpression lines. **F-J,** τ_cl_, Pn, chlorophyll content, tillering and yield per plant in WYG7 and PIF4^v3m^ rice line. The parameter τ_cl_ represents half-time of stomatal closure speed calculated according to Vico et al., 2011. Each bar data represents the mean of replicates (*n*=7) ±SE for panel **A**, and *n*=10 for panels **B-I**. One-way *ANOVA* was used to determine the significance level between WYG7 and *PIF4*-OE lines, while *t*-test was used to determine the significant levels between WYG7 and PIF4^v3m^.

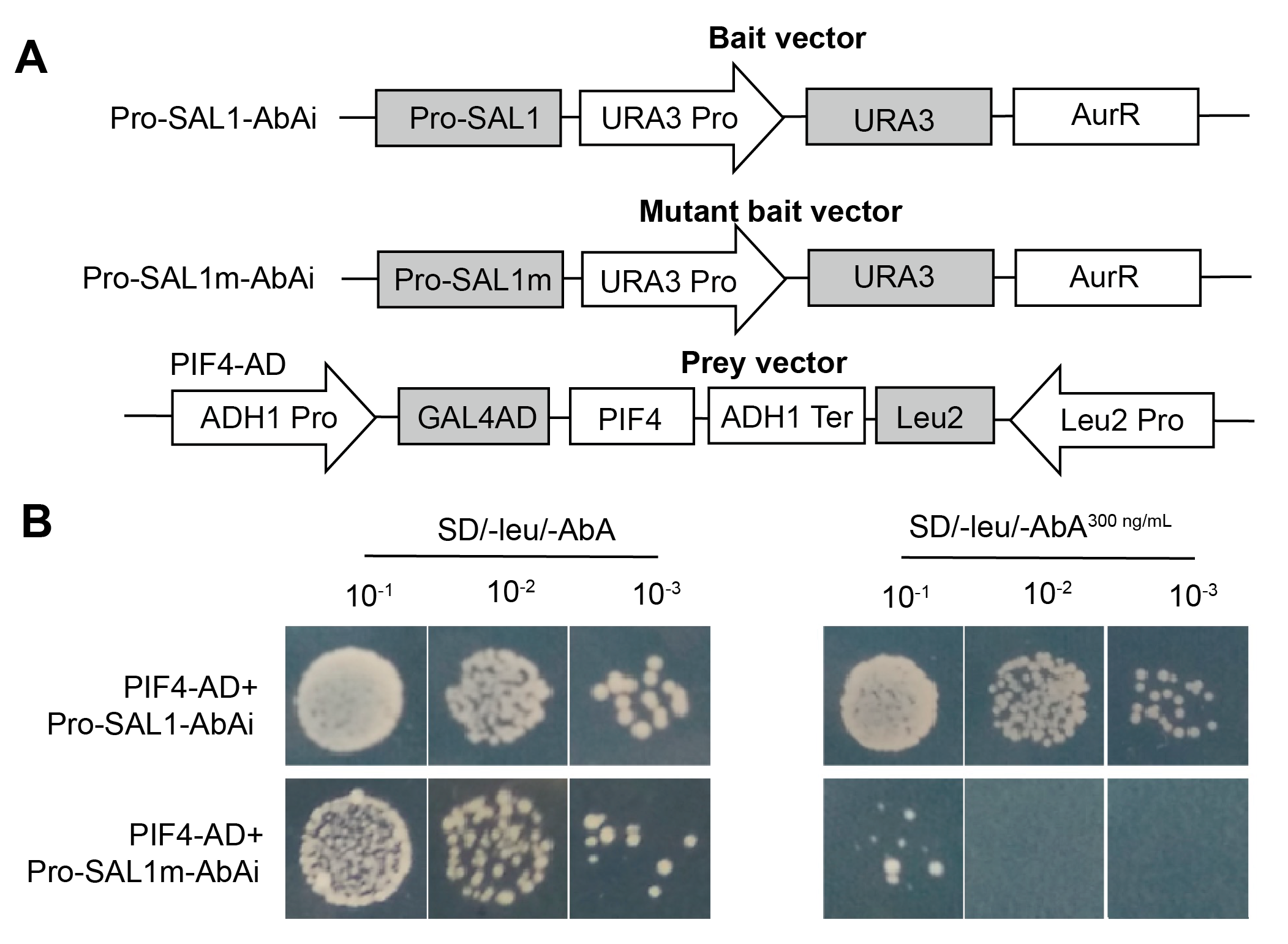

**Figure S8.** PIF4 specifically binds to G-box motif present in *SAL1* promoter. **A**, Schematic diagrams of the promoter and mutant promoter fragments in *SAL1* used for the construction of the bait and mutant bait vectors. **B**, Y1H assays showing the interaction between PIF4 and the CACGTG-motif present in the *SAL1* promoters, based on the ability of the transformed yeast strains to grow on SD/−Leu/AbA^300ng/mL^ medium with gradient dilution (1/10, 1/100, 1/1000). The transformants grown on SD/−Leu/−AbA plate were used as positive controls for transformants growth. Positive transformants were confirmed by spotting yeast cells onto agar medium of SD/−Leu with 300 ng/mL AbA. These assays were repeated three times with similar results.

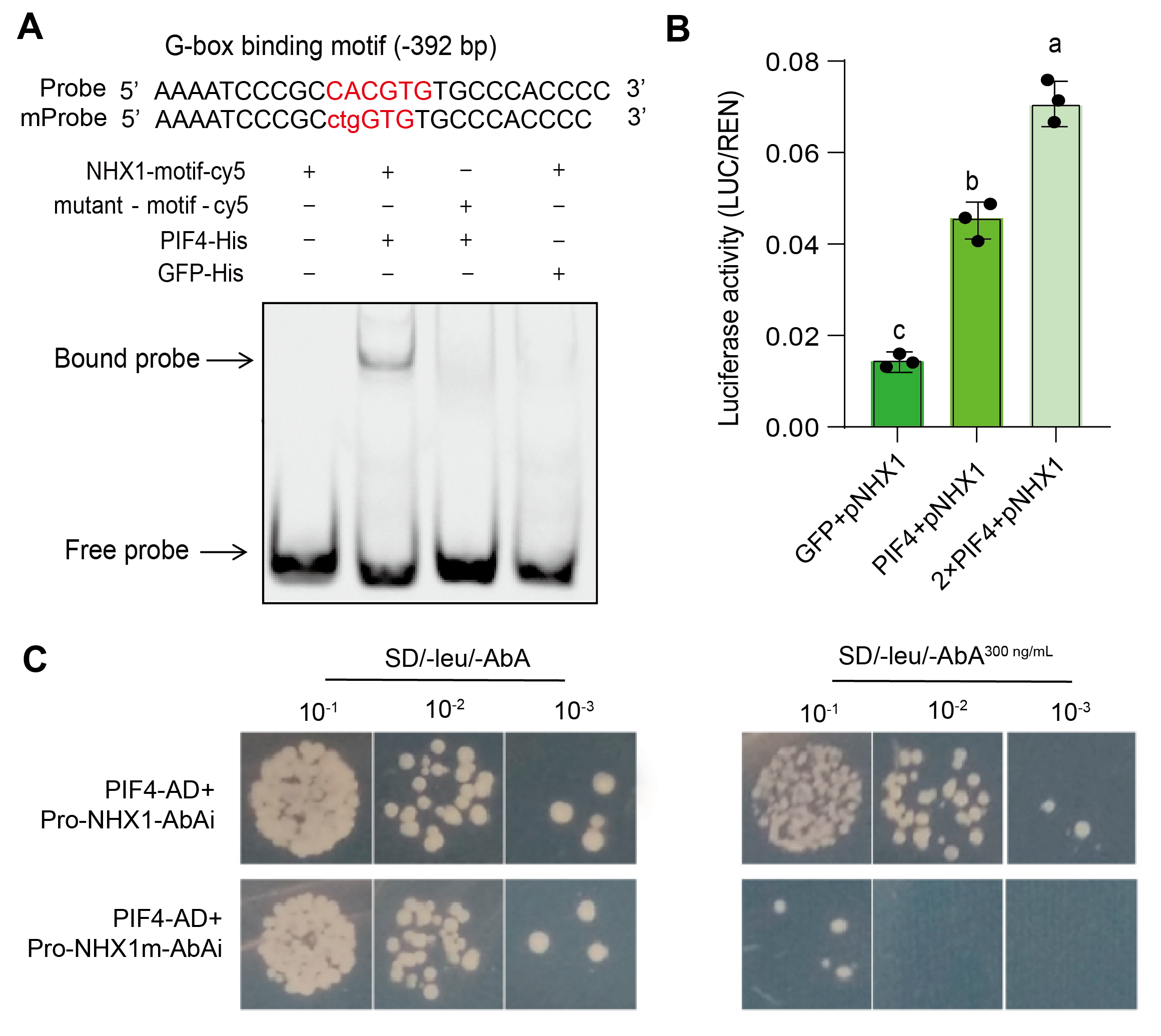

**Figure S9.** PIF4 specifically binds to G-box motif of *NHX1* promoter. **A**, EMSA experiments representing PIF4 binds to G-box motif present in *NHX1* promoter. **B**, luciferase activity determination for *PIF4* transcriptionally activates the expression of *NHX1*. *n*=3 for panels **B. C**, Y1H assays showing the interaction between OsPIF4 and the CACGTG-motif present in the *NHX1* promoters.

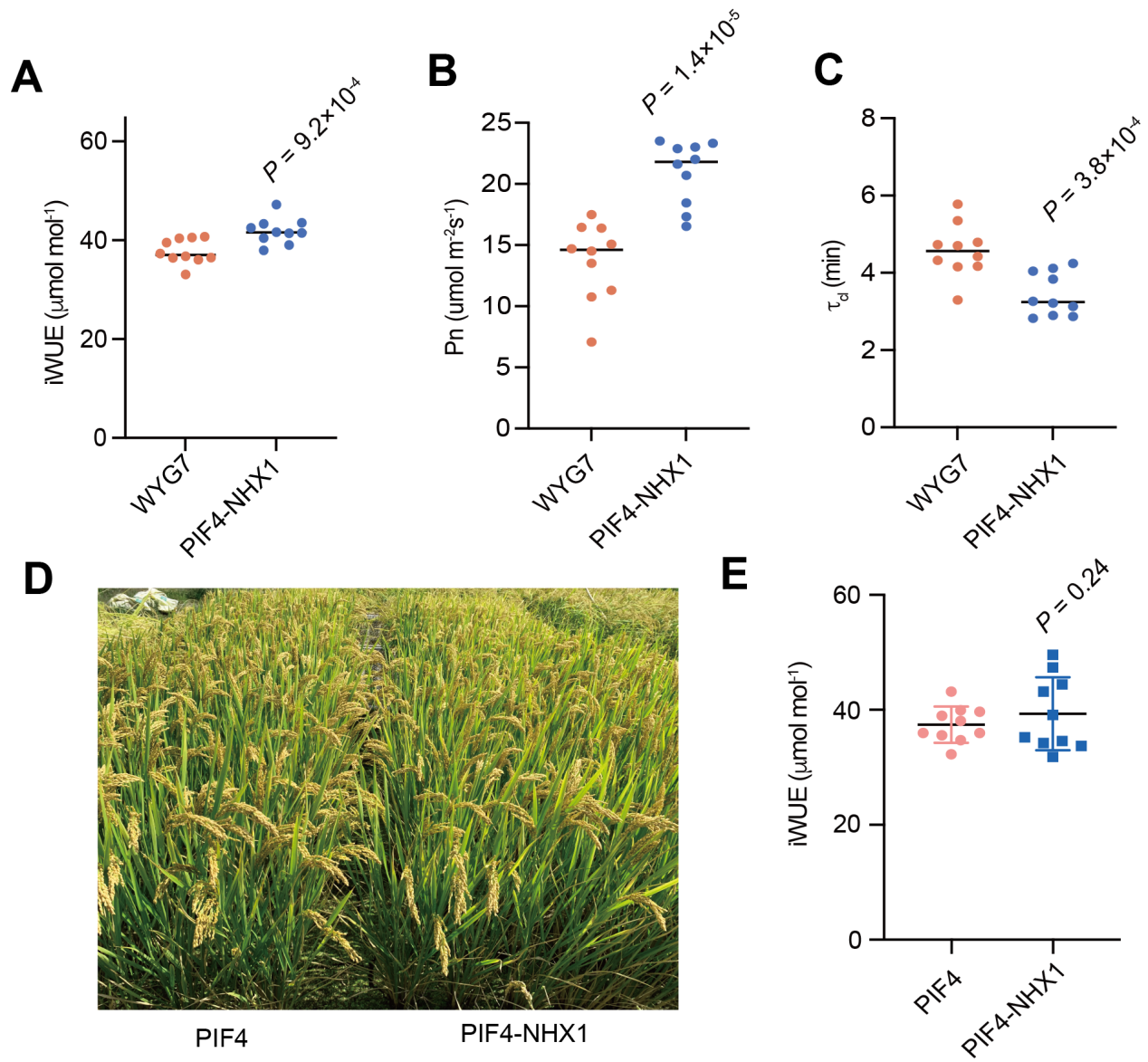

**Figure S10.** Comparison on iWUE and Pn between WYG7 and co-expression of *PIF4* and *NHX1* (*PIF4*-*NHX1*) at graining stage. **A**, iWUE; **B**, Pn. **C**, stomatal closure speed (τ_cl_). The τ_cl_ represents half-time of stomatal closure during high light switched to low light according to Vico et al., 2011. **D**, field performance between *PIF4*-OE and co-overexpression lines of *PIF4*-*NHX1* under DS at graining stage. **E**, iWUE_FL_ between *PIF4*-OE and co-overexpression lines of *PIF4*-*NHX1*. Each bar data represents the mean of replicates (*n*=10) ±SE. One-way *ANOVA* was used to determine the significance level between WYG7 and *PIF4*-*NHX1* and between *PIF4*-OE and *PIF4*-*NHX1*.
